# Clu1/Clu form mitochondria-associated granules upon metabolic transitions and regulate mitochondrial protein translation via ribosome interactions

**DOI:** 10.1101/2024.08.17.608283

**Authors:** Leonor Miller-Fleming, Wing Hei Au, Laura Raik, Pedro Guiomar, Jasper Schmitz, Ha Yoon Cho, Aron Czako, Alexander J Whitworth

**Affiliations:** MRC Mitochondrial Biology Unit, University of Cambridge, Cambridge Biomedical Campus, Cambridge, CB2 0XY, United Kingdom

## Abstract

Mitochondria perform essential metabolic functions and respond rapidly to changes in metabolic and stress conditions. As the majority of mitochondrial proteins are nuclear-encoded, intricate post-transcriptional regulation is crucial to enable mitochondria to adapt to changing cellular demands. The eukaryotic Clustered mitochondria protein family has emerged as an important regulator of mitochondrial function during metabolic shifts. Here, we show that the *Drosophila melanogaster* and *Saccharomyces cerevisiae* Clu/Clu1 proteins form dynamic, membraneless granules adjacent to mitochondria in response to metabolic changes. Yeast Clu1 regulates the translation of a subset of nuclear-encoded mitochondrial protein by interacting with their mRNAs while these are engaged in translation. We further show that Clu1 regulates translation by interacting with polysomes, independently of whether it is in a diffuse or granular state. Our results demonstrate remarkable functional conservation with other members of the Clustered mitochondria protein family and suggests that Clu/Clu1 granules isolate and concentrate ribosomes engaged in translating their mRNA targets, thus, integrating metabolic signals with the regulation of mitochondrial protein synthesis.

## INTRODUCTION

Mitochondria perform many essential cellular functions, most notably in the production of ATP and regulation of cell death [1, 2]. To perform these myriad functions, mitochondria are extremely dynamic organelles, trafficking along the cytoskeletal tracks, undergoing fission and fusion events, and interacting with other organelles [3]. This dynamic nature allows mitochondria to respond rapidly to changes in cellular conditions such as transient stresses or fluctuations in nutrient availability [4]. Consequently, disruption of mitochondrial function causes wide-ranging cellular defects and human diseases [5].

While mitochondria contain their own genome and translation machinery, the majority of their proteome is nuclear encoded, transported to mitochondria, imported and sorted to their appropriate destination [6]. The central role of various mitochondrial activities in various cellular and metabolic processes means that their protein composition is also highly dynamic [7, 8]. Thus, transcriptional and translational changes must be tightly regulated and coordinated between nuclear and mitochondrial genomes, particularly in response to acute changes in metabolic demands. An essential layer of regulation occurs at the post-transcriptional level, where RNA-binding proteins play a crucial role [9, 10]. These proteins are essential for the stability, localization, and translation of mRNAs, and their dysregulation can underlie different diseases [10].

The Clustered mitochondria protein family is a conserved group of proteins found among eukaryotes, which includes the yeast Clustered mitochondria (Clu1) [11], the *Drosophila* Clueless (Clu) [12] and the mammalian Clustered mitochondria homologue (CLUH) [13], which share a common mitochondrial clustering phenotype and mitochondrial dysfunction when mutated [11–14]. Knockout mice for *Cluh* are neonatal lethal, while fly mutants for *Clu* mostly die during development with rare adult flies only surviving a few days [12, 15, 16], highlighting the critical roles these proteins play in different organisms during crucial developmental and metabolic transitions.

CLUH is an RNA-binding protein that preferentially binds and regulates mRNAs of multiple nuclear-encoded mitochondrial proteins involved in the tricarboxylic acid cycle (TCA) cycle, oxidative phosphorylation (OXPHOS) and several other mitochondrial metabolic pathways [13]. However, how this regulation is achieved is still unclear. Emerging evidence has shown that these proteins form foci under different metabolic conditions suggesting these structures are membraneless organelles which may play an important role into how the Clustered mitochondria proteins function [17, 18].

Here, we exploited the tractability of *Drosophila* and budding yeast as model systems and investigated in detail the cellular and molecular dynamics of Clu/Clu1 granules, their nature and composition, analysed the yeast Clu1 binding partners *in vivo*, and explored the relationship of Clu1 with mRNA translation. We propose a model where Clu/Clu1 granules isolate ribosomes engaged in translating their targets in response to metabolic changes to regulate their translation.

## RESULTS

### Clu/Clu1 forms dynamic foci upon metabolic transitions in flies and yeast

Clu was recently reported to have a dynamic localization in *Drosophila* egg chambers, affected by nutritional status [18]. We independently observed that a line expressing GFP-tagged endogenous Clu (GFP-Clu) exhibited a punctate subcellular distribution in various fly tissues (Figure S1A). Notably, we observed that Clu localization was indeed dynamic in *Drosophila* egg chambers (Figure 1A). After fasting overnight (16 h), GFP-Clu was diffuse in the cytoplasm of nurse cells and follicle cells (Figure 1A). When flies were refed, GFP-Clu rapidly formed distinct foci within 30 min, which gradually increased in size over time (Figure 1A). After 6 h of refeeding, these puncta reached remarkable sizes, particularly in the nurse cells, with an average area of 2.37 µm^2^ (up to 40 µm^2^) (Figure 1A, Figure S1B-E). Interestingly, when flies were ‘re-fasted’, Clu rapidly returned to the diffuse state within minutes (Figure 1A). These results show that Clu localization is dynamic and regulated by metabolism.

**Figure 1.**
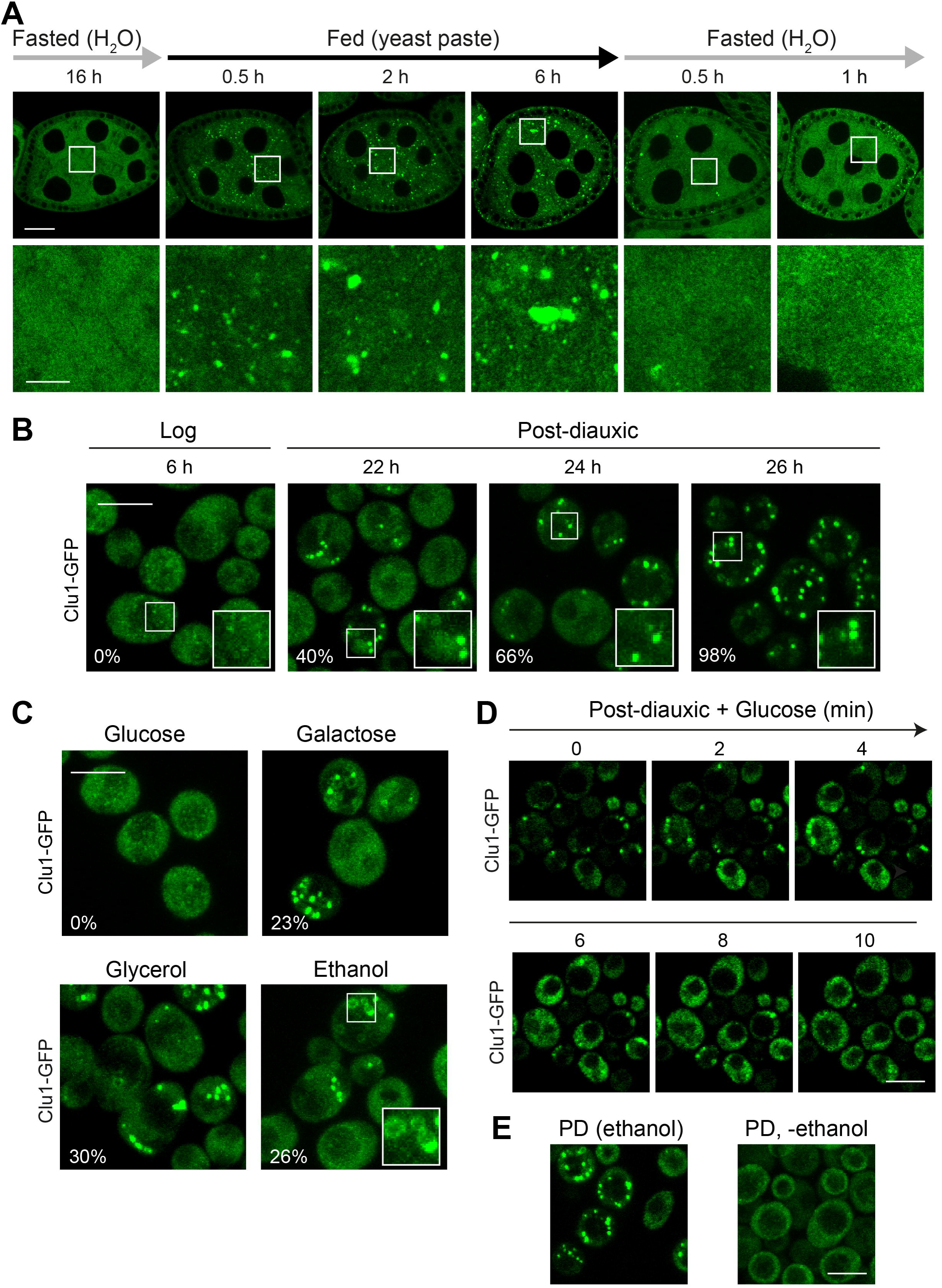
*Drosophila* Clu and yeast Clu1 form dynamic foci. (A) Confocal imaging of egg chambers isolated from GFP-Clu flies. One-day-old female flies were mated and fed with yeast paste for 2 days, fasted overnight (16 h), refed for 6 h, and re-fasted for 1 h. Images represent the indicated time points during the time-course and are representative of at least three biological replicates. (B) BY4741 yeast cells expressing endogenously tagged Clu1-GFP were grown in media containing glucose and analysed by confocal imaging in exponential (log) and PD (PD) phases. White box regions show Clu1 foci magnified (inset). Percentage of cells showing large foci is indicated in each image (>200 cells counted). (C) Mid-log phase cells grown in media containing glucose were washed and incubated with media containing galactose, glycerol or ethanol and analysed by confocal microscopy after 6 h. White box region is magnified (inset) showing Clu1-GFP in a ring like pattern. (D) PD Clu1-GFP cells were layered on an agarose pad made with media containing 2% glucose and imaged over a 10 min time course. (E) PD Clu1-GFP cells were washed and incubated in SC media without any carbon source for 20 min. Cells were imaged and compared with cells kept in the original media. Scale bars: A (top), 20 μm; A (bottom), 5 μm; B-E, 5 μm.

*Saccharomyces cerevisiae* is very well defined metabolically, and the metabolic state can easily be manipulated by changing nutrient availability in the media. Thus, we assessed whether the yeast Clu orthologue, Clu1, which to date is poorly characterised, also showed a dynamic localisation. For this, we used a GFP-tagged endogenous *CLU1* strain (Clu1-GFP) grown under different metabolic conditions. In the logarithmic (log) phase, characterised by exponential growth and fermentative metabolism, Clu1-GFP was mainly diffuse in the cytoplasm with some cells showing some undefined small puncta (Figure 1B, and Figure S1F), as previously reported [18]. Upon transition to the post-diauxic phase (PD), when glucose is exhausted and cells undergo a metabolic transition into respiration (Figure S1F), Clu1-GFP progressively formed very bright, distinct foci such that nearly all cells (98%) had Clu1-GFP foci 10 h after the diauxic shift (Figure 1B, and Figure S1F). This suggested that Clu1 forms foci when metabolism becomes reliant on mitochondrial respiration.

To confirm whether the transition from fermentation to respiration triggered Clu1 foci formation, we switched cells grown in glucose media to media containing respiratory carbon sources such as ethanol, glycerol and galactose. Indeed, these abrupt metabolic transitions also led to the formation of Clu1-GFP foci, sometimes appearing as rings (Figure 1C), supporting that the transition to respiration is the trigger for the formation of foci.

To understand whether this phenomenon was reversible, similar to what we observed in *Drosophila* egg chambers, we added back glucose to PD cells. Within minutes, Clu1-GFP foci dissipated and became diffuse throughout the cytoplasm (Figure 1D, Video S1). In addition, we also found that Clu1 foci dissipate upon removing the carbon source (ethanol) from the media of PD cultures (Figure 1E), suggesting that foci require ATP to be maintained. Together, these results demonstrate that Clu1 localisation is extremely dynamic in yeast, forming slowly upon the metabolic transition from fermentation to respiration, and disassembling rapidly upon reverting their metabolism back to fermentation.

### Clu1 foci are distinct from PBs and SGs and form upon mitochondrial stress

Stress granules (SGs) and Processing bodies (P bodies; PBs) are cytoplasmic membraneless organelles, also known as biomolecular condensates, composed of RNAs and proteins. SG formation is induced, while PBs become more abundant, under different stresses including nutritional stress [19, 20]. As PBs and SGs form in the PD and stationary phases respectively [21, 22], and are quickly reversed by the addition of glucose, we questioned whether Clu1 was a component of these structures.

First, we tested whether the Clu1 foci appeared upon stresses known to trigger SGs and/or PBs: heat shock, carbon starvation, hypotonic stress, hyperosmotic stress, oxidative stress, and mitochondrial stress (sodium azide) [19, 20, 22, 23]. In contrast to SGs and PBs, Clu1 foci were not formed in most tested conditions (Figure 2A). Interestingly, treatment with sodium azide, a complex IV inhibitor, rapidly induced formation of distinct Clu1-GFP foci, resembling the foci observed upon the transition to respiration (Figure 2A). Curiously, another mitochondrial stressor, carbonyl cyanide m-chlorophenyl hydrazone (CCCP), a protonophore that dissipates the mitochondrial membrane potential, did not trigger formation of Clu1 foci (Figure 2A).

**Figure 2.**
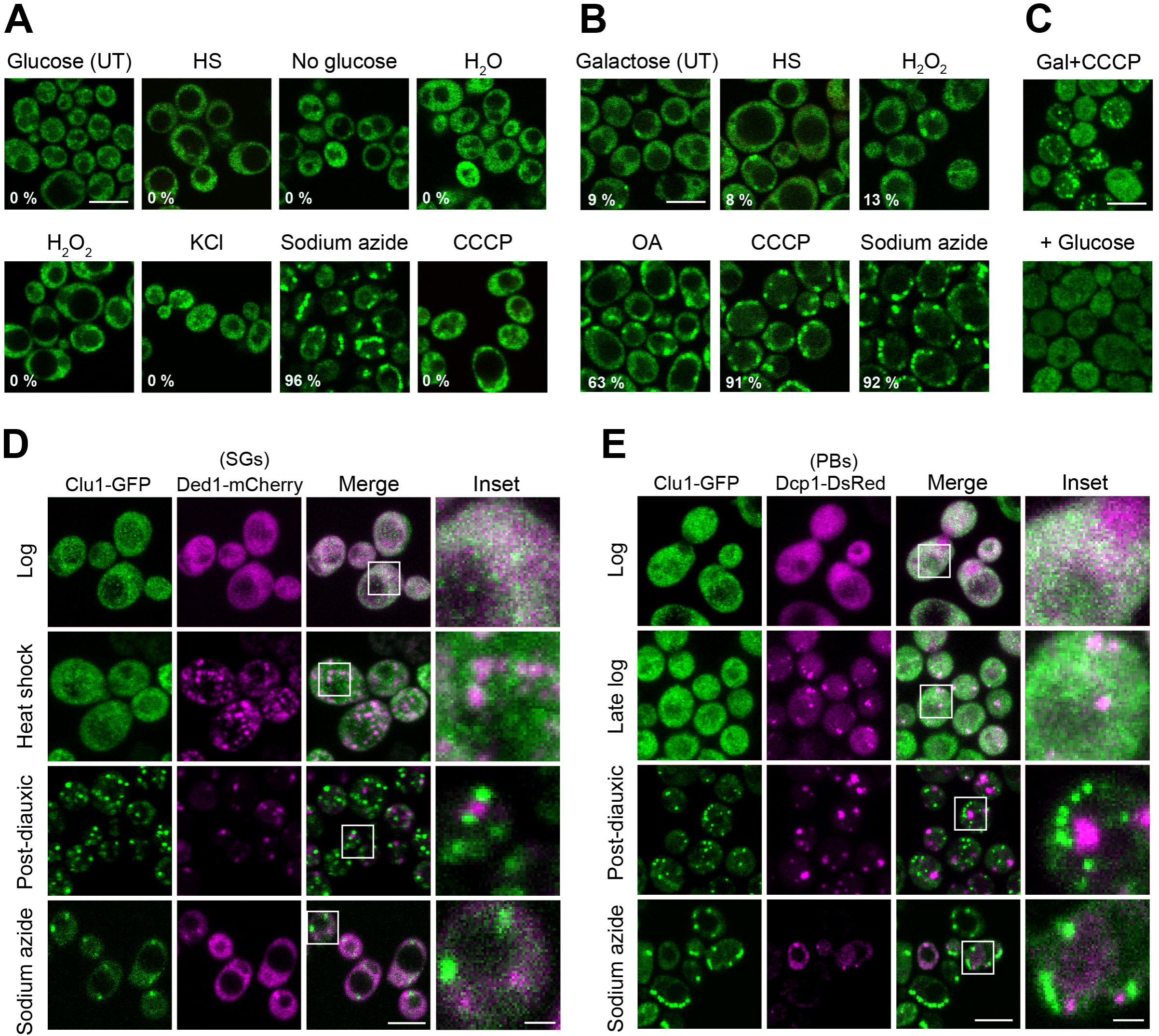
Yeast Clu1 form foci under mitochondrial stresses and do not co-localise with PBs and SGs. (A) Maximum intensity projection of confocal images of Clu1-GFP cells grown in glucose-containing media until mid-log phase, then subjected to different stresses: heat shock (HS; 46 °C, 30 min), without any carbon source (No glucose, 10 min), exposure to hyperosmotic stress (1 M KCl, 30 min), hypotonic stress (H_2_O, 10 min), oxidative stress (3 mM H2O2, 15 min), and mitochondrial stress sodium azide (0.5%, 15 min) and CCCP (30 μM, 15 min), or untreated (UT). The percentage of cells containing Clu1 foci is indicated in each image. (B) Confocal imaging (maximum intensity projection) of mid-log Clu1-GFP cells shifted from glucose to galactose-containing-media and incubated for 6 h, then subjected to stresses as in A, plus oligomycin (10 µM) and antimycin A (40 µM) (OA; 30 min). The percentage of cells containing Clu1 foci is indicated in each image. (C) Maximum intensity projection of Clu1-GFP cells grown in galactose-containing media for 6 h, treated with CCCP for 15 min (Gal+CCCP), spiked with glucose and imaged 20 min later (+ Glucose). (D) Confocal microscopy of Clu1-GFP strain expressing Ded1-mCherry, a marker of stress granules (SGs) in the log and PD phases, and after heat shock or sodium azide treatment, as described above. (E) Confocal microscopy images of Clu1-GFP strain expressing Dcp1-DsRed, a marker of P-bodies (PBs) in the log, late log and PD phases, and after sodium azide treatment. Scale bars = 5 µm; D, E inset = 2 µm.

We also investigated the localisation behaviour of Clu1 when cells were more reliant on respiration. For this, we took Clu1-GFP cell cultures that had been shifted from glucose-to galactose-containing media, in which foci started to be formed, and subjected them to heat shock, oxidative stress or mitochondrial stressors. Similar to fermenting conditions, neither heat shock nor oxidative stress triggered the formation of Clu1 foci. In contrast to what was described for Clu foci in flies [18], Clu1 foci are not sensitive to oxidative stress as this treatment did not lead to their disappearance (Figure 2B). Interestingly, treatments with the mitochondrial stressors sodium azide, CCCP or oligomycin and antimycin (inhibitors of mitochondrial ATP synthase and complex III, respectively) enhanced the formation of Clu1 foci in all cells (Figure 2B). Spiking glucose into respiring cells grown with galactose and treated with CCCP led to the disappearance of Clu1 foci (Figure 2C), showing the foci are also reversible following mitochondrial stress.

SGs and PBs vary their composition according to the type of stress that leads to their formation [19, 20, 24]. Thus, although yeast Clu1 foci did not form under most typical stress conditions that trigger SGs and PBs, this did not exclude the possibility that Clu1 could be a component of these structures specifically during metabolic shift or mitochondrial stress. To test this hypothesis, we co-expressed Clu1-GFP with Ded1-mCherry, a marker of SGs [25], or Dcp1-DsRed, a marker of PBs [26]. We analysed whether these markers co-localise under conditions where both Clu1 foci and SGs or PBs are formed, specifically during PD phase and after treatment with sodium azide. None of these conditions led to the co-localisation of Clu1 foci with these SG or PB markers (Figure 2D, E). It is worth noting that the number of cells with PBs started increasing in late log phase as previously described [21], while Clu1 foci only appeared later in the PD phase (Figure 2E). This supports that Clu1 is not a component of SGs or PBs.

In the fly egg chambers, GFP-Clu foci also did not colocalise with SG or PB markers, Fmr1 immunostaining and Tral-mRFP, respectively (Figure S2), as previously reported [18]. Thus, our results show that despite some similarities between Clu1 foci and SGs and PBs, yeast and *Drosophila* Clu1/Clu foci are different structures.

### Clu foci are biomolecular condensates adjacent to mitochondria

While analysing yeast Clu1 localisation in PD cells, we consistently observed Clu1 foci localised near mitochondria (Figure 3B-D), and even appeared between adjacent mitochondria, likely coinciding with recent fission events (Figure 3B-D). In a few cells, Clu1 appeared in a ring shape surrounding a mitochondrion (Figure S3A). We observed a similar distribution of Clu1 foci formed under respiratory stress (cells shifted to galactose and treated with CCCP), indicating that a different trigger of foci formation leads to a similar behaviour (Figure 3E). In addition, Clu1 foci partially co-localised with foci of the mitochondrial fission factors, Mdv1 (Figure 3F) and its paralog Caf4 (Figure S3B) [27]. The close association of Clu/Clu1 foci with mitochondria and their response to changing metabolic conditions is consistent with Clu/Clu1 performing an important role in maintaining mitochondrial function. Indeed, while knockout of *CLU1* in yeast (*clu1*Δ) has no impact on growth in glucose-containing media (Figure S3C), viability and growth were affected in ethanol-containing media, when respiration is essential (Figure S3D), as previously reported [18]. Consistent with this, we observed that mitochondrial morphology is only clustered under respiratory conditions (Figure S3E, F).

**Figure 3.**
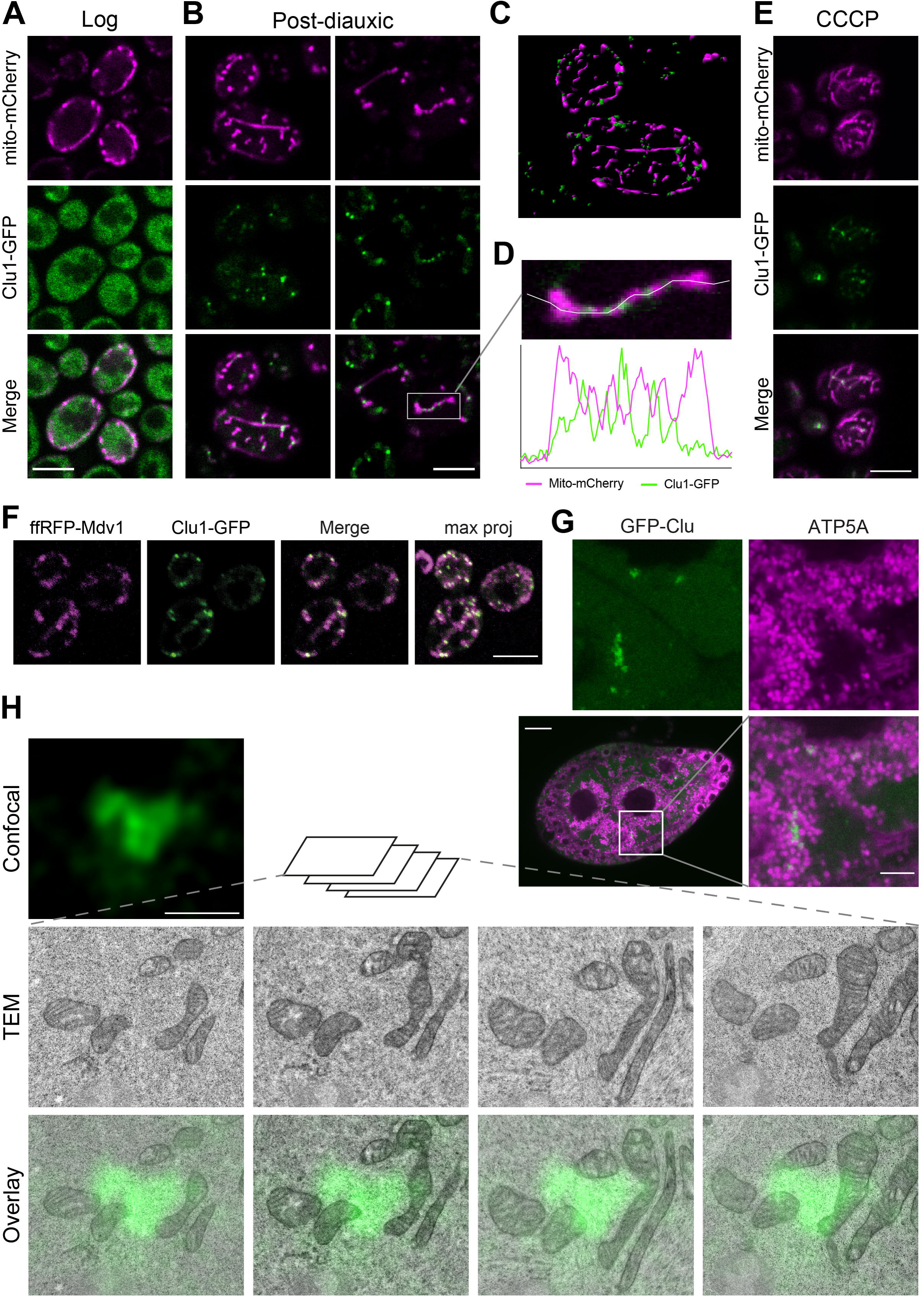
Clu1/Clu foci are localized in close proximity to mitochondria in yeast and flies. (A, B) Confocal microscopy of yeast Clu1-GFP cells co-expressing the mitochondrial marker mito-mCherry in the log (A) and PD phases (B). (C) 3D rendering of cells in PD phase. (D) Inset of panel B with an intensity profile plot showing the fluorescence distribution of Clu1-GFP and mito-mCherry. (E) Confocal image of Clu1-GFP cells expressing mito-mCherry after being shifted to galactose for 6 h and treated with CCCP. (F) Confocal imaging of Clu1-GFP cells co-expressing mitochondrial fission marker ffRFP-Mdv1 grown into PD phase. (G) Confocal imaging of a GFP-Clu egg chamber from 3-day-old flies (6 h of feeding after overnight fasting) immunostained for ATP5A to visualise mitochondria. (H) Correlative light and electron microscopy images of a GFP-Clu egg chamber nurse cell showing the juxtaposition of a GFP-Clu granule and mitochondria. The serial transmission electron microscopy (TEM) images, taken at 5000 × magnification, correspond to the confocal image shown. Scale bars: A, B, E, F = 5 µm; G = 20 µm, inset = 5 µm; H = 1 µm.

In *Drosophila*, Clu foci were localised near mitochondria (Figure 3G), as previously described [12]. To further investigate the subcellular localisation of Clu/Clu1 foci, we employed correlative light and electron microscopy (CLEM). In ‘refed’ *Drosophila* egg chambers, where GFP-Clu foci are abundant, we found that Clu foci consistently localized adjacent to mitochondria, not directly overlapping (Figure 3H). Notably, we did not observe any organelle or distinct structure between the Clu foci and mitochondria, suggesting that they are likely in direct contact. Moreover, GFP-Clu foci did not appear to be bound by any detectable membrane, suggesting they form a membraneless organelle (Figure 3H).

The biophysical properties of membraneless organelles have been intensely characterised recently and have been found to form by liquid-liquid phase separation (LLPS) driven by weak and multivalent interactions [28]. To explore whether Clu/Clu1 foci are formed by LLPS, we treated ovaries from fed flies and PD yeast cells, which exhibit numerous Clu/Clu1 foci, with 1,6-hexanediol, an aliphatic alcohol which disassembles LLPS condensates by disrupting weak hydrophobic interactions [29–32]. Live imaging of GFP-Clu in egg chambers treated with 1,6-hexanediol showed a steady dissolution of the GFP-Clu foci over time in contrast to untreated egg chambers (Figure 4A). Complementing this, Clu1-GFP yeast cells were permeabilised with digitonin and incubated with either 5 or 10 % 1,6-hexanediol. With 5% incubation, both the number and size of Clu1 foci decreased while with 10%, the foci were completely dissolved (Figure 4B). Taken together, the preceding results suggest that Clu/Clu1 foci have membraneless nature formed by LLPS, thus, hereafter, we refer to them as Clu/Clu1 granules.

**Figure 4.**
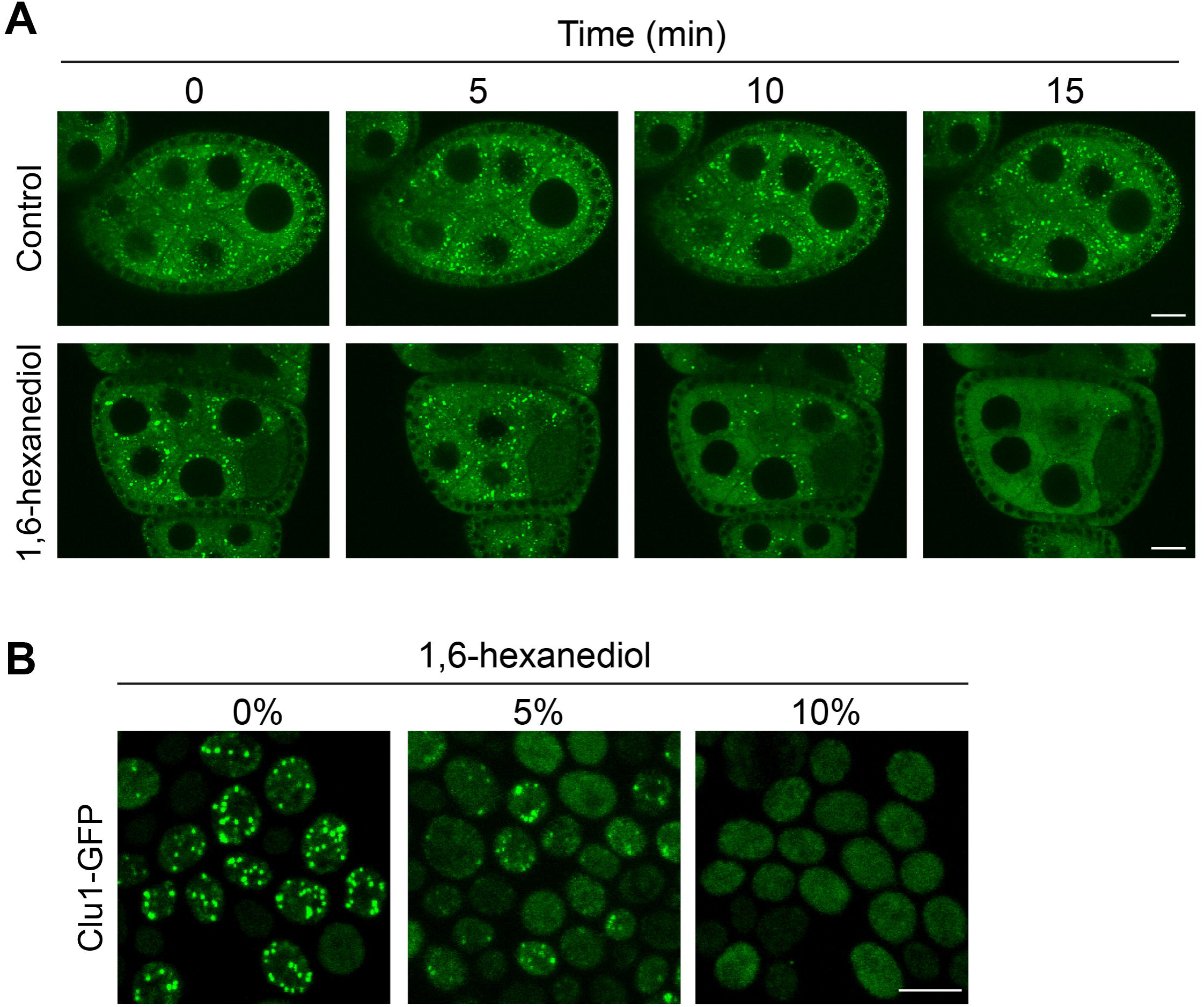
Clu/Clu1 granules have liquid-like properties. (A) Representative live imaging of a GFP-Clu egg chamber incubated in Schneider’s media containing insulin, treated with 1,6-hexanediol or vehicle and imaged over a 15 min period. (B) Images of PD Clu1-GFP cells fixed after being permeabilised with digitonin for 2 min, followed by 5 min treatment with 0, 5 and 10% 1,6-hexanediol. Scale bars = 20 µm (A) and 5 µm (B).

### Clu granules contain RNA and require mRNAs engaged in translation

Many biomolecular condensates that contain RNA-binding proteins rely on RNAs for their formation [33–35]. Evidence supports that Clu/Clu1 are RNA-binding proteins, similar to CLUH, as they co-purify with mRNAs [16]. Thus, we hypothesised that Clu granules contain RNAs which may influence Clu granule stability. To test this, cultured egg chambers from refed flies were treated with RNase A which led to the complete disassembly of most Clu granules (Figure 5A, B), similar to the effect on PBs [36]. This indicates that RNA plays a crucial role in the formation and/or stability of Clu granules.

**Figure 5.**
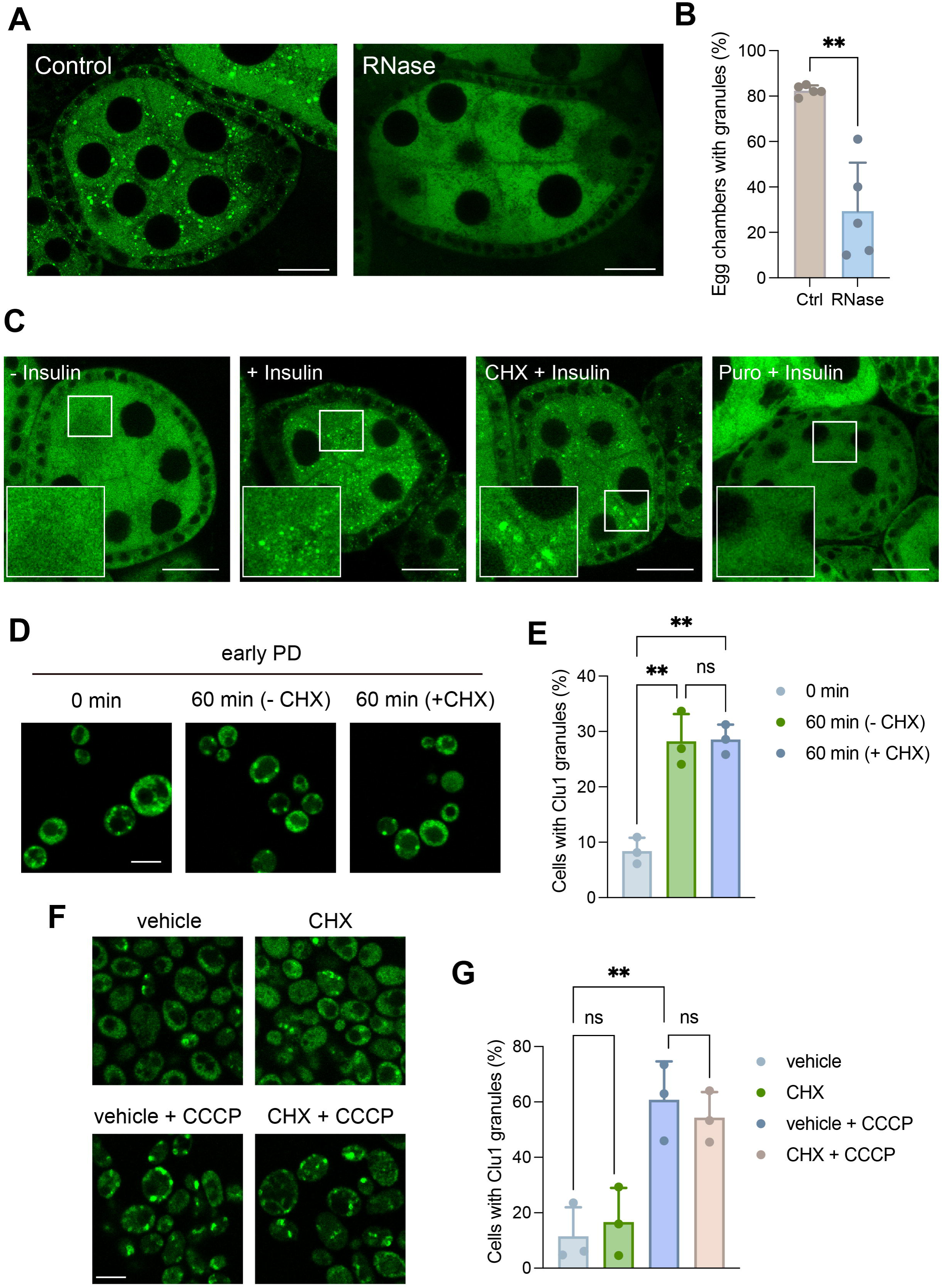
Clu/Clu1 granules are RNase- and puromycin-sensitive but cycloheximide-insensitive. (A) Dissected ovaries from GFP-Clu flies permeabilized and incubated with or without RNase for 5 min, fixed and analysed by confocal microscopy. (B) Quantification of the percentage of egg chambers containing Clu granules per fly shown in A (mean ± SD; unpaired t-test with Welch’s correction; n = 5 animals; ** P < 0.01). (C) Confocal images of egg chambers subjected to different treatments after being dissected in Schneider’s media: a control with no additional treatment (40 min) (-Insulin), incubation with insulin (30 mins) 10 min after dissection (+ Insulin), or pre-treatment with CHX (CHX + Insulin) or puromycin (Puro + Insulin) for 10 min followed by incubation with insulin (30 mins). (D) Confocal images and (E) quantification of Clu1-GFP cells in the early PD phase and treated with or without CHX for 60 min. (F) Confocal images and (G) quantification of Clu1-GFP cells with granules in cells shifted to galactose-containing media for 6 h, pre-treated with CHX or vehicle for 10 min and then followed by CCCP or vehicle treatment for 15 min. Data (E and G) are presented as mean ± SD; n = 3 biological replicates; > 100 cells each biological replicate (one-way ANOVA, ns = non-significant, ** P < 0.01). Scale bars = 20 µm (A, C) and 5 µm (D, F).

Since the formation of SGs and PBs depends on mRNAs released from ribosomes, we sought to determine whether the formation of *Drosophila* Clu granules similarly depends on a similar mechanism. In yeast, flies and mammalian cells, pre-treatment with cycloheximide (CHX) – an inhibitor of protein synthesis that traps mRNAs in polysomes – prevents the formation of SGs and PBs upon stress exposure [19, 36–38]. To test this, we took advantage of the fact that insulin can trigger granule formation in dissected egg chambers (Video S2) [18]. Contrary to SGs and PBs, pre-treatment with CHX did not prevent the formation of Clu granules upon addition of insulin (Figure 5C), indicating that Clu granules do not form with mRNAs that exit translation. However, this did not exclude the possibility that Clu granules may form with free mRNAs that have not yet engaged in translation. Therefore, we next assessed the effect of puromycin, which dissociates ribosomes from the mRNA by releasing truncated polypeptides [39]. In contrast to CHX, pre-treatment with puromycin before insulin addition, prevented the formation of Clu granules (Figure 5C), indicating that Clu granule formation depends on mRNAs being actively engaged in the ribosome.

To investigate whether yeast Clu1 behaves like fly Clu, we incubated Clu1-GFP cells in early PD phase with CHX and quantified Clu1 granule formation. Similar to our observation in fly egg chambers, the addition of CHX did not affect the formation of Clu1 granules compared to the control (Figure 5D, E). Notably, pre-treatment with CHX also did not prevent the formation of Clu1 foci in mitochondrial stress conditions (respiring cells treated with CCCP) (Figure 5F, G), showing again that Clu1 granules formed during a gradual shift to respiration or by acute mitochondrial stress are similar in nature.

Taken together, these results indicate that Clu/Clu1 granule formation depend on mRNAs being engaged in translation, but not on the release of mRNAs from ribosomes, highlighting a unique aspect of their assembly mechanism compared to other RNA-containing granules like SGs and PBs.

### Yeast Clu1 regulates the translation of nuclear-encoded mitochondrial proteins

To further characterize the role of Clu1 in yeast, we performed BioID, a proximity-dependent biotinylation assay in vivo [40]. This assay uses BirA*, a promiscuous variant of the biotin ligase BirA that labels neighbouring proteins [41], allowing us to identify potential direct or indirect interactors, as well as proteins localised near Clu1.

We generated a Clu1-GFP-BirA* strain by integrating BirA* 3’ to Clu1-GFP in the genome and verified that this did not affect the dynamic localisation of Clu1 (Figure S4). As a control, we generated a strain with cytosolic BirA*. This identified 26 proteins in proximity to Clu1 in the log phase and 22 in the PD, 4 of which are common between the two phases (Figure 6A, and Table S1). Notably, 12 proteins detected in the log phase are nuclear-encoded mitochondrial proteins, most of which localise in the mitochondrial matrix (Figure 6A, and Table S1). Additionally, we detected three cytosolic ribosomal proteins and the translation initiation factor eIF-4B (Tif3). Most of the mitochondrial proteins identified are involved in metabolic processes. These include Pda1 and Lat1, components of the pyruvate dehydrogenase complex that converts pyruvate into acetyl-CoA; Aconitase 1 (Aco1) a participant in the TCA cycle; and the mitochondrial aldehyde dehydrogenases Ald4 and Ald5, which are involved in acetate production. In the PD phase, we identified fewer nuclear-encoded mitochondrial proteins, with Aco1 and Lat1, a component of the pyruvate dehydrogenase complex being common to both phases; five ribosomal proteins, with RpL20a common to the log phase, and the translation initiation factor eIF5 (Tif5). The RNA-binding protein Scp160 and the tRNA synthetase Ses1 were also identified, which are encoded by genes highly co-expressed with Clu1 according to the SPELL database [42] with correlation scores of 3.6 (the most correlated gene) and 3.1, respectively.

**Figure 6.**
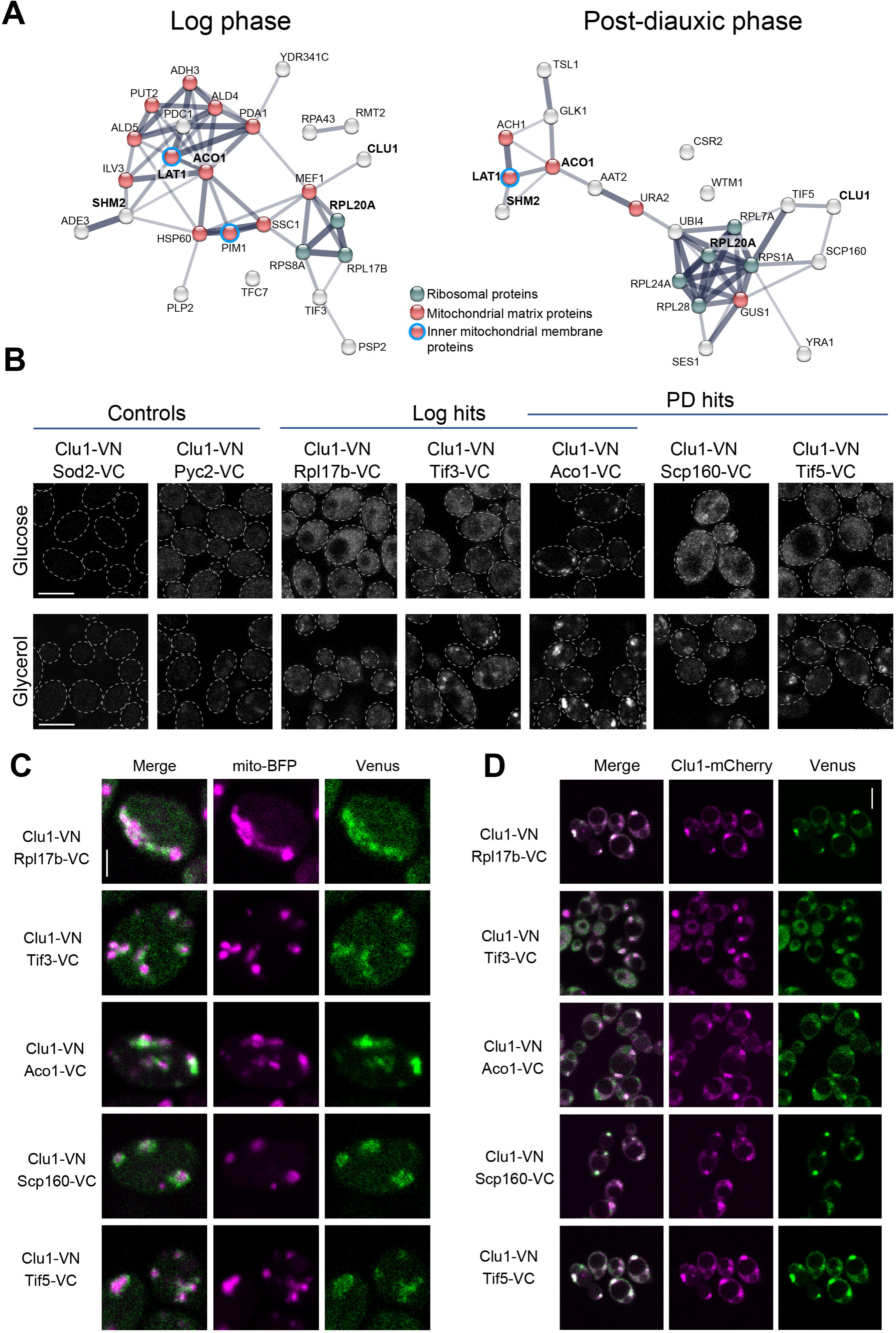
Clu1 is in a complex with ribosomes and mitochondrial proteins. (A) STRING association network analysis (physical and functional interactions based) of yeast Clu1 interactors identified by BioID in the log or PD (PD) phases. Highlighted in bold are proteins found in common in both growth phases. (B) Confocal microscopy analysis of bimolecular fluorescent complementation (BiFC) assay of cells co-expressing Clu1 fused with the Venus N-terminal fragment (Clu1-VN) and either non-interactor controls (Sod2 and Pyc2) or BioID hits (Rpl17b, Tif3, Aco1, Scp160, and Tif5) fused to the Venus C-terminal fragment (protein*-*VC). Cells were either grown in glucose media until mid-log phase (upper panels) or shifted to glycerol media for 24 h (lower panels). (C, D) Localisation of where Clu1-BioID hits interact visualised by the Venus reconstitution (green) in relation to mitochondrial marker mito-BFP (magenta) (C) and Clu1-mCherry (magenta) (D) in cells shifted to glycerol for 24 h. Scale bars = 5 (B, D) and 2 µm (C).

To validate the protein-protein interactions identified by BioID *in vivo*, we used the bimolecular fluorescence complementation assay (BiFC) [43, 44]. We tested strains co-expressing Clu1 tagged with the N-terminal fragment of the fluorescent protein Venus (VN) [45] and selected BioID hits (Rpl17b, Aco1, Scp160, Tif3 and Tif5) and two control non-interactors (Sod2 and Pyc2), each fused with the C-terminal fragment of Venus (VC) at their respective C-termini [46]. The split-Venus constructs were integrated into the yeast genome to maintain physiological expression levels [45, 46]. After growing the different strains in fermenting conditions (glucose) and shifted to respiring conditions (glycerol), we assessed the reconstitution of Venus fluorescence by confocal microscopy. This confirmed the interaction of Clu1 with the BioID hits in both carbon sources, and, as expected, no signal was observed in the controls (Figure 6B). In glucose, the Venus signal was generally diffuse, except for cells co-expressing Clu1-VN and Aco1-VC, which displayed bright puncta (Figure 6B). In glycerol, we observed a change in Venus localization from diffuse to punctate in cells co-expressing Clu1-VN and Rpl17b-VC, Scp160-VC, Tif3-VC, or Tif5-VC. Additionally, there was an increase in the size and number of puncta in the Clu1-VN Aco1-VC strain (Figure 6B). To understand the localisation of these puncta we co-expressed these strains with a mitochondrial marker (mito-BFP) or endogenously tagged Clu1-mCherry. In all cases, we observed that the reconstituted Venus puncta localised close to the mitochondria (Figure 6C) and co-localised with Clu1-mCherry granules in respiratory conditions (Figure 6D), confirming that indeed these interactions happen in granules. These results indicate that Clu1 co-localises with cytosolic ribosomal and nuclear-encoded mitochondrial proteins, likely just before these mitochondrial proteins are imported into the mitochondria.

The proximity of Clu1 to ribosomal and nuclear-encoded mitochondrial proteins led us to investigate whether Clu1 regulates translation by analysing the translatome of *CLU1* knockout cells compared to the wild-type strain. To achieve this, we used puromycin-associated nascent chain proteomics (Punch-P), which consists of isolating intact polysomes from cells that will incorporate biotinylated puromycin into the nascent polypeptides giving a snapshot of translation at the time of harvesting. These biotinylated polypeptides can then be purified using streptavidin and identified by mass spectrometry (Figure 7A) [47, 48].

**Figure 7.**
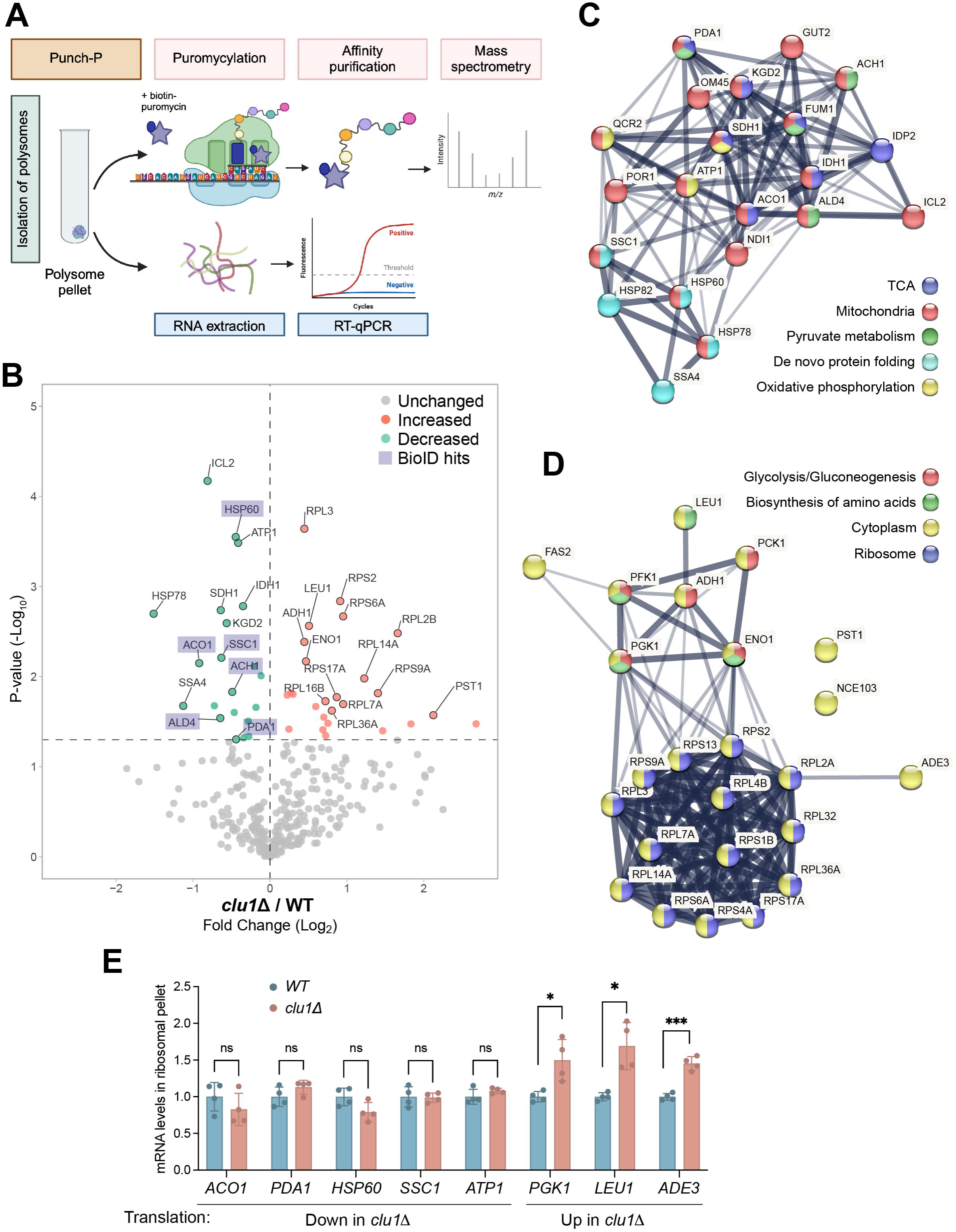
Translation of specific nuclear-encoded mitochondrial proteins is affected by the absence of Clu1. (A) Experimental workflow schematic of the Punch-P assay. Punch-P involves incorporating biotinylated-puromycin into newly synthetised proteins via polysomes followed by purification and identification of the newly synthetised proteins by mass spectrometry. Polysomes isolated by a sucrose cushion were used either for the Punch-P assay, or isolation of RNA and qRT-PCR. (B) Volcano plot of the proteins detected in the Punch-P assay comparing control with *clu1*Δ cells. The X-axis represents the fold change in translation of proteins and the Y-axis corresponds to the p-value. Shaded hits correspond to proteins found in proximity to Clu1 (BioID hits). (C, D) STRING interaction network of the proteins less translated (C) or more translated (D) in the *clu1*Δ compared to control. (E) qRT-PCR analysis of mRNA of selected hits that are decreased or increased in the Punch-P assay in control or *clu1*Δ cells normalised by the 25S rRNA levels (mean ± SD; unpaired t test; n = 4 samples; * P < 0.05, *** P < 0.001). Illustrations were created using BioRender.

We detected 360 proteins, 47 of which were significantly different between wild-type and *clu1*Δ strains (Figure 7B, and Table S2). Of these 47 proteins, 21 were less translated in the absence of Clu1 (Figure 7B). Notably, except Hsp82, Ssa4 and Idp2, all these proteins are nuclear-encoded mitochondrial proteins. Although Hsp82 is not a mitochondrial protein, it functions closely with Tom70 targeting proteins into the mitochondria [49]. The majority of proteins that were less translated in the absence of Clu1 are metabolic proteins involved in the TCA cycle, pyruvate metabolism and oxidative phosphorylation. The remaining proteins are chaperones (Figure 7C). Notably, 6 out of the 21 proteins (Figure 7B), such as Aco1 and Pda1, were also identified in the BioID assay as being in proximity to Clu1 (Figure 6A), further supporting the idea that Clu1 regulates the translation of these proteins. The proteins that were more translated in the absence of *CLU1* were all cytosolic proteins and included ribosomal and metabolic proteins involved in glycolysis/gluconeogenesis or amino acid biosynthesis (Figure 7D).

To investigate whether the reduced signal from the Punch-P assay could be due to reduced abundance of those transcripts, we extracted RNA from the same polysomes used for the Punch-P and quantified transcripts that were differentially translated. Surprisingly, we found no differences in the levels of mRNAs that were less translated in the *clu1*Δ strain (*ACO1*, *PDA1*, *HSP60*, *SSC1* and *ATP1*) (Figure 7E). However, we observed higher levels of mRNAs encoding the more translated proteins (*PGK1*, *LEU1* and *ADE3*). These results strengthen the hypothesis that Clu1 regulates the efficiency of translation of specific nuclear-encoded mitochondrial mRNAs, while the increased mRNAs/proteins may be upregulated as a compensatory mechanism.

### Clu1 interacts with nuclear-encoded mitochondrial mRNAs, via ribosome interaction

Our findings suggested that yeast Clu1 regulates the translation of nuclear-encoded mitochondrial mRNAs, similar to the mammalian orthologue CLUH [13]. Since CLUH is known to directly bind to mRNAs as an RNA-binding protein [13], we explored whether Clu1 functions in a similar way. We further tested the hypothesis that Clu1 interacts with mRNAs while they are being translated. For this, we performed RNA immunopurification (RIP) of Clu1-GFP and GFP (control) under conditions where ribosomes were kept assembled and stalled by CHX, and compared with conditions where ribosomes were dissociated by EDTA or puromycin. We predicted that transcripts whose translation was dysregulated in the absence of Clu1 (Punch-P assay) and/or encode proteins found in its proximity (BioID assay) interact physically with Clu1. Thus, we selected *ACO1, PDA1, HSP60, SSC1, ATP1* and *ADH3* to quantify by RT-qPCR. As negative controls, we selected transcripts encoding two nuclear-encoded mitochondrial proteins not predicted to interact with Clu1 (*SOD2* and *ATP2*) and two non-mitochondrial proteins (*TAF10* and *UBC6*). The mRNAs that we predicted to interact with Clu1 were highly enriched in the Clu1-GFP RIP, in contrast to the negative controls, when ribosomes were intact (Figure 8A). Strikingly, these transcripts were no longer enriched when ribosomes were dissociated with EDTA or puromycin (Figure 8B, C). In parallel, we quantified the rRNA, and found both 18S and 25S rRNA enriched in the Clu1-GFP RIP when ribosomes were intact. This enrichment was also lost upon dissociation of ribosomes (Figure 8D). These results support that Clu1 forms a complex with ribosomes, and that Clu1 is interacting with specific nuclear-encoded mitochondrial mRNAs only when they are engaged in translation. We further corroborated the association of Clu1 with the ribosome by co-immunoprecipitating Clu1 and the ribosomal protein Rpl3 (Figure 8E).

**Figure 8.**
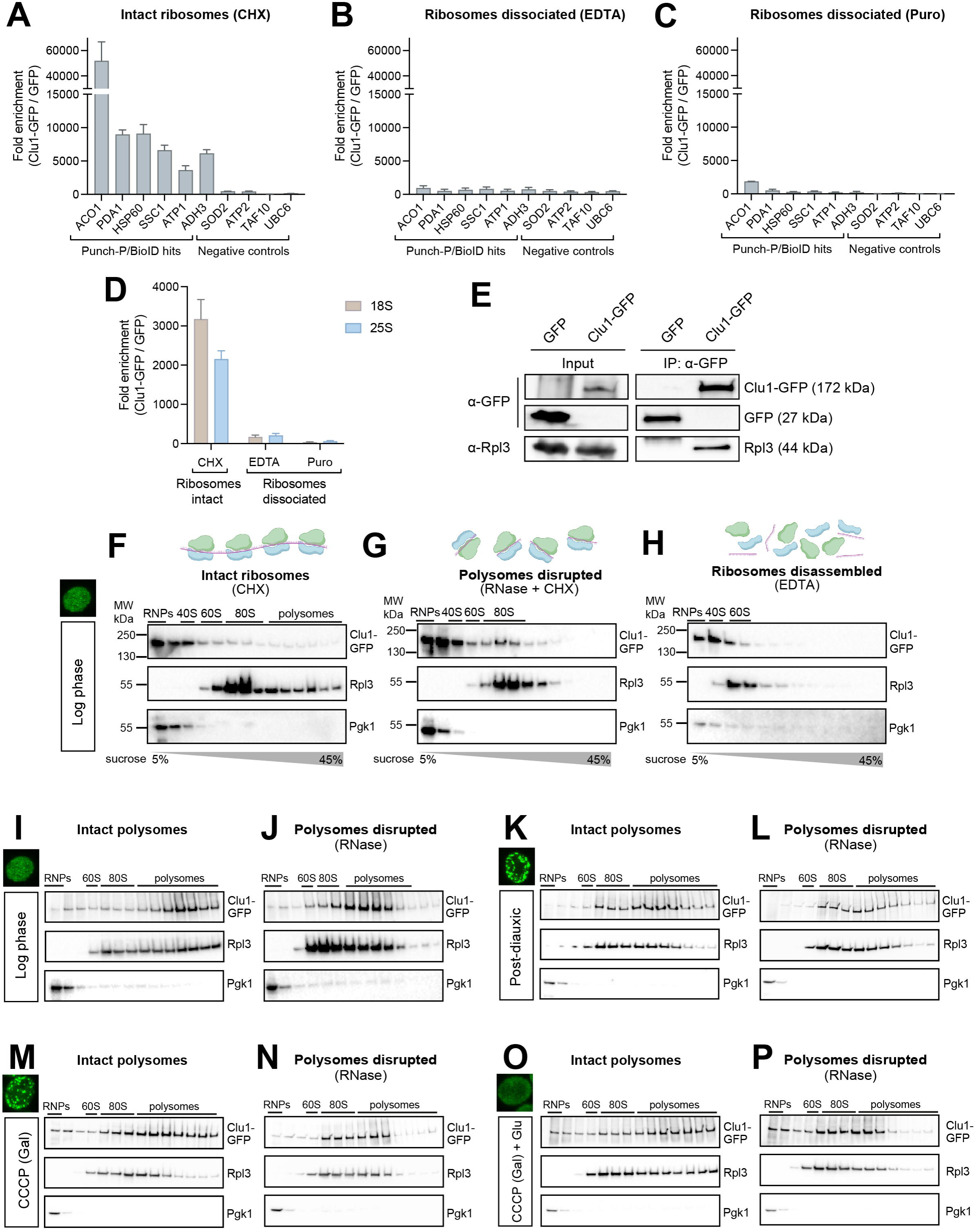
Clu1 interacts with translating nuclear-encoded mitochondrial mRNAs. (A-C) Clu1-GFP RNA immunoprecipitation (RIP) was performed under ribosome-preserving (CHX) or ribosome-disassembling conditions (EDTA or puromycin (Puro)), using GFP-expressing cells as a control. RNA was extracted following IP of Clu1 and mRNA levels of BioID and/or Punch-P hits (proteins less translated in *clu1*Δ) (*ACO1*, *PDA1*, *HSP60, SSC1, ATP1* and *ADH3*) were quantified by qRT-PCR. The levels of *ATP2*, *SOD2, TAF10* and *UBC6* mRNAs were also quantified as non-interactor controls. mRNA levels detected in RIP were normalised by input levels. Following normalization, the Clu1 RIP levels were divided by the control RIP levels. (mean ± SD, n = 3) (D) qRT-PCR analysis of RIP samples as in A-C for 18S and 25S rRNA. (E) Immunoblot of GFP IP control and Clu1-GFP IP, detecting physical interaction with the ribosomal protein Rpl3. (F-H) Sucrose continuous density gradients and ribosome profiling performed under ribosome-preserving (CHX), polysome disrupting (RNase+CHX) or ribosome-disassembling (EDTA) conditions, analysing the co-sedimentation of Clu1-GFP with ribosomes (Rpl3) by immunoblot of log phase grown cells. Pgk1 serves as an indicator of cytosolic/soluble protein sedimentation. (n = 3) (I-P) Sucrose density gradients and ribosome profiling, similar to F-H, performed following *in vivo* formaldehyde cross-linking to preserve interactions under different growth conditions: log phase (I, J) or PD phase (K, L); or metabolic/stress conditions: CCCP (gal; galactose) (M, N) or CCCP (gal) ± glucose (O, P). (J, L, N, P) Sucrose gradients of same lysates used for I, K, M, O treated with RNase to disrupt polysomes. Illustrations were created using BioRender.

To further explore the association of Clu1 with ribosomes and its potential changes under varying metabolic conditions, we performed sucrose gradient fractionation and assessed its co-migration with ribosomes. In addition, we questioned whether the Clu1 interaction with the ribosome was mRNA dependent. For this, we analysed cells in the log phase under three different conditions: with CHX to maintain monosomes and polysomes integrity; with CHX followed by RNase treatment that converts polysomes into monosomes; and with EDTA to disassemble polysomes and monosomes. The ribosome profile was monitored by 254 nm absorbance (not shown), by immunoblotting for the large ribosomal subunit Rpl3, and as a control, we immunoblotted for Pgk1, a glycolytic monomeric enzyme not involved in ribosomal or high-order protein complexes [50]. When polysomes and monosomes were preserved (CHX), Clu1 was abundant in the light fractions, corresponding to free Clu1 or Clu1 binding to small proteins or RNAs, but was also present in the high-density fractions co-migrating with monosomes and polysomes (Figure 8F). Upon treating the lysate with RNase, as expected, polysomes were disrupted and ribosomes accumulated in the monosome fractions. Here, Clu1 disappeared from the heaviest fractions and became enriched in the monosome fractions (Figure 8G), indicating that the interaction with ribosomes is mRNA independent. Upon disassembling ribosomes with EDTA, Clu1 completely disappeared from the high-density fractions, as did ribosomes, further confirming its association with the ribosome (Figure 8H). The similarity between Clu1 and ribosome migration profiles in all conditions tested in the log phase indicates that Clu1 binds ribosomes, either in polysome or monosome form, independently of the interaction with mRNA.

To further study Clu1 in the context of ribosomes, we chemically cross-linked Clu1-GFP log cells to stabilize weaker interactions that might dissociate during the long sucrose gradient ultracentrifugation procedure. Cross-linking significantly shifted the migration pattern of Clu1, from the lighter fractions to now being highly enriched in the polysome fractions (Figure 8I), indicating that the interaction between Clu1 and ribosomes may be transient and/or weak. To confirm that the presence of Clu1 in the polysome fraction is due to its interaction with the ribosome, we treated the cross-linked lysate with RNase (Figure 8J). Although this treatment had a modest effect in disrupting polysomes, Clu1 clearly followed the ribosomes’ migration pattern, indicating that Clu1 migrates into the denser fractions of the gradient via an interaction with ribosomes.

To determine whether the association of Clu1 with the ribosome changes upon metabolic transitions or mitochondrial stress when Clu1 is in granules, we also analysed cross-linked cells in PD phase or under respiratory stress (Galactose + CCCP). In both conditions, the abundance of high density polysomes decreased indicating a reduced general translation, and notably, although some Clu1 shifted towards the monosome fractions, it was still highly abundant in polysomes fractions (Figure 8K, M). We further assessed the Clu1–ribosome interaction when granules disappear upon shifting the metabolism towards fermentation, as we observed when spiking glucose into the media of cells grown in respiratory media and then stressed with CCCP (Galactose + CCCP) (see Figure 2C). Upon brief (15 min) incubation with glucose, both ribosomes and Clu1 moved to the heavy polysomes fractions (Figure 8O) when compared to cells before glucose addition (Figure 8M). Altogether, these results show that Clu1 is interacting with polysomes independently of the metabolic condition and its localisation.

As previously, to confirm that this co-migration is due to an mRNA-independent association of Clu1 with ribosomes, we treated these lysates with RNase and observed that the Clu1 migration profile completely mirrored the ribosome profile in all conditions tested (Figure 8L, N, P). Altogether, these results show that the majority of Clu1 is always associated with ribosomes, independently of the mRNA interaction, whether Clu1 is diffuse or in granules, and independently of whether the metabolic shift is gradual or acute.

## Discussion

Mitochondria are key organelles for cellular adaptation to metabolic shifts and stress. Post-transcriptional regulation of their proteome plays a crucial role in how mitochondria respond to various conditions. The *Drosophila* protein Clu and its mammalian orthologue CLUH were shown to be involved in regulating translation and maintaining mitochondrial function [11–17, 51–54], but their precise roles are poorly understood.

In this study, we provide novel insights into the function of the *Drosophila* Clu and the yeast orthologue Clu1. Both proteins dynamically change their localization upon metabolic transitions, shifting from a diffuse cytosolic distribution to granule formations adjacent to mitochondria, suggesting a potential regulatory mechanism in mitochondrial metabolism. The dynamic distribution of fly Clu triggered by fasting and refeeding or by the addition of insulin to cultured egg chambers corroborates earlier observations [18]. Egg chambers are cellular structures highly sensitive to nutritional changes [55], however, the exact metabolic changes that occur during these rapid nutritional fluctuations, or insulin addition, are poorly characterised. Nevertheless, Clu granules appear in a transition from low to high metabolic conditions, which likely translates into higher respiratory metabolism [56]. In yeast, we precisely defined the metabolic trigger for the formation of Clu1 granules as the transition from fermentation to respiration, a process involving profound transcriptional, translational and metabolic changes, particularly at the mitochondria level [57]. Our findings are in line with a recent study demonstrating that CLUH forms granules in primary mouse hepatocytes when cells are switched from nutrient-rich to nutrient-poor media, indicating that granule formation in response to metabolic changes is a conserved feature [17].

Membraneless organelles are cellular compartments that, despite lacking a lipid membrane, play important roles in organizing the cellular content. When dysregulated, they may contribute to the pathogenesis of diseases such as amyotrophic lateral sclerosis [35, 58]. These organelles are formed through liquid-liquid phase separation (LLPS), involving specific proteins and, in many cases, mRNAs and is driven by weak and transient interactions between multivalent molecules [28, 58]. Their rapid assembly and disassembly along with the easy exchange of components with the external environment allows cells to quickly adapt to metabolic changes and various stresses. Previous studies had observed that Clu1/Clu/CLUH proteins form foci, but definitive evidence that these are biomolecular condensates, including confirming the absence of a membrane, was lacking [17, 18]. Despite several similarities with PBs and SGs, such as being rapidly dynamic and reversible, Clu1/Clu granules are distinct structures, not co-localising with classic markers of these organelles and not forming under most typical stress conditions. This shows functional similarities to the mammalian CLUH granules, which were also shown to be independent of SGs despite co-localising with the typical SG component G3BP1 [17]. Using a CLEM approach, we found that fly Clu granules are not delimited by a membrane and are in close proximity to mitochondria. In addition, we observed that Clu/Clu1 granules quickly dissolve upon treatment with 1,6-hexanediol which, despite some caveats related to its use [59, 60], is consistent with these granules being formed by LLPS [29–32].

Building on recent observations of Clu/CLUH dynamic localisation, we focussed our investigation on the nature and formation of the Clu/Clu1 granules, including their components. We first found that Clu granules depend on RNA for stability, as treatment of egg chambers with an RNase (where live imaging with this treatment is feasible) resulted in the rapid disappearance of Clu granules. Interestingly, Clu granule formation was prevented by dissociating ribosomes with puromycin, indicating that granules require mRNAs engaged in translation to be formed. Corroborating our results, a study in primary mouse hepatocytes showed that a subset of CLUH granules incorporated puromycin when ribosome dissociation was prevented with emetine, supporting that at least a subset of CLUH granules contain translating ribosomes [17]. In contrast, we found that stalling translation with CHX had no impact on granule formation in flies or yeast, consistent with observations for CLUH in HeLa cells [17] and in contrast to SGs and PBs [19, 36–38]. Altogether, these results indicate that Clu/Clu1 granule formation is tightly linked to mRNAs engaged in translation.

Our data strongly implicates a role for yeast Clu1 in regulating the translation of specific nuclear-encoded mitochondrial mRNAs. First, the characterisation of Clu1 interactome by BioID showed mainly ribosomal proteins, nuclear-encoded mitochondrial proteins, RNA-binding proteins and translation initiation factors. In line with our findings, a recent BioID analysis of CLUH from a human colorectal carcinoma cell line revealed a similar interactome [61]. In addition, Clu1, Clu, CLUH and the *Arabidopsis thaliana* orthologue, Friendly mitochondria (FMT) [62], were shown to interact with ribosomal proteins, highlighting the conservation of this interaction mechanism [16, 51]. Interestingly, characterising the interaction of Clu1 with a subset of the interacting proteins (Scp160, Tif3, Tif5, Rpl17, and Aco1) by BiFC, we found that these interactions occur regardless of Clu1 condensation state and metabolic state, suggesting that Clu1 role is independent of its condensation state.

Our translatome approach further supports the role of Clu1 in regulating the translation of specific mitochondrial proteins. We found that the absence of Clu1 decreases the efficiency of translation of various nuclear-encoded mitochondrial proteins, many of which we identified as being in proximity to Clu1 by BioID. It remains to be determined whether this decrease in translation efficiency is caused by a reduced number of ribosomes per polysome (indicating possible decrease in translation initiation) or by slower elongation rates.

Additionally, we found that mRNAs encoding proteins that we found proximal to Clu1 and/or dysregulated in its absence, such as *ACO1, PDA1* and *HSP60*, physically interact with Clu1. Notably, the orthologous mammalian mRNAs were also shown to interact with CLUH [13], revealing a remarkable conservation of the protein-RNA interaction and functional homology between these proteins.

Surprisingly, our Clu1 RIP findings show that Clu1 interacts with mRNAs only when monosomes and polysomes are intact, supporting that Clu1 only interacts with mRNAs that are engaged in translation. Furthermore, our sucrose gradients showed that Clu1 associates with polysomes and monosomes independently of Clu1 interacting with mRNA. These results indicate that Clu1 interaction with mRNAs depends on translating ribosomes and involve a direct interaction with the ribosome. The striking similarity between the interacting mRNAs found in yeast and human cells implies that Clu1 and CLUH have a conserved function. As CLUH was shown to directly bind mRNAs [13, 51, 61] and to interact with ribosomal proteins [51, 61], we propose that Clu1 and CLUH interact simultaneously with the ribosome and mRNAs, with this interaction occurs specifically when mRNAs are being translated, likely by influencing their localization, stability, or translation efficiency.

Importantly, we found that Clu1 associates the ribosomes when they are in polysomes, likely while actively translating, in both its diffuse state and when it forms granules, whether triggered by a switch to respiration or mitochondrial stress. The prevention of Clu granule formation by dissociating ribosomes with puromycin suggests that the presence of ribosomes is crucial for this process. Altogether, these findings support the idea that Clu1/Clu granules contain translating ribosomes. Hence, since membraneless organelles can facilitate or limit interactions of their components, by concentrating interacting molecules or isolating specific macromolecules from the surrounding [63], we hypothesise that Clu/Clu1 granules may promote or pause the translation of these proteins even in adverse conditions where perhaps the mRNA targets of Clu1/Clu would be degraded.

Further work is needed to fully elucidate the complexity of these mechanisms, but our findings help define Clu/Clu1 granules as unique membraneless organelles that likely regulate the production of mitochondrial proteins in proximity to the mitochondria, which has important implications for understanding the dynamic and homeostatic processes regulating mitochondrial function. It will be interesting to investigate in the future if the unique subcellular environment within the granules has an impact in the regulation of translation of these transcripts.

## MATERIAL AND METHODS

### *Drosophila* methods Stocks and husbandry

*Drosophila melanogaster* flies were raised under standard conditions in a temperature-controlled incubator with a 12 h:12 h light:dark cycle at 25 °C and 65% relative humidity, on food consisting of malt extract, molasses, cornmeal, yeast, agar, soya powder, water, propionic acid, nipagin. GFP-Clu trap line (clu[CA06604]) was obtained from Rachel Cox and Tral-RFP was kindly provided by Daniel St Johnston [64].

### Fed and refed time course

Crosses were set up with 20 GFP-Clu females and 10 males (one-day-old flies) in bottles with yeast paste for one day. Flies were starved overnight in tubes with tissue paper wet in water only. Flies were refed with yeast paste containing bromophenol blue (to control for feeding) up to 6 h and then starved again in wet paper up to 1 h and a half. Female ovaries were dissected in different intervals on Grace’s media (Thermo Fisher Scientific; 11605045) when flies were being fed or PBS when starved and then fixed in 4% formaldehyde (Thermo Fisher Scientific; 28908) made in PBS containing 0.3% of Triton X-100 (PBS-T) for 15 mins at room temperature with rotation. After fixation, ovaries were washed in PBS-T for 10 min 3 times. Next, ovarioles were carefully separated, mounted in Vectashield Antifade Mounting Medium (Vector Laboratories, Inc.; H-1000) and imaged on a Zeiss LSM880 confocal microscope (Carl Zeiss MicroImaging) with Nikon Plan-Apochromat 63x/1.40 NA oil immersion objective.

### Immunostaining

For Fmr1 immunostaining, one-day-old GFP-Clu female flies were crossed with males in a ratio of 3:1 and fed yeast paste for two days. Flies were then starved overnight with tissue paper wet in water (starved flies) and refed with yeast paste for 2 h (refed flies). For ATP5A immunostaining, 1-day-old flies were fed in yeast paste for 2 days, fasted overnight and refed for 6 h. Ovaries were dissected in Grace’s media (Thermo Fisher Scientific; 11605045), fixed in 4% PFA made in 0.3% TritonX-100 PBS (PBS-T) for 15 min at room temperature with rotation and washed three times, 10 min with PBS-T. Subsequently, they were blocked in 1% BSA + PBS-T for 1 h, incubated with an antibody against Fmr1 (DSHB Hybridoma; 5A11) in 0.3% PBS-T containing 1% BSA, diluted 1:50 or an antibody against ATP5A (Sigma; ab14748), used at 1:500 dilution, overnight at 4 ^°^C. Following the incubation with primary antibody, ovaries were washed three times, 10 min with PBS-T and incubated with the goat anti-mouse secondary antibody, Alexa Fluor™ 594 (Invitrogen; A11005) at 1:250 dilution made in 1% BSA PBS-T in the dark for 1 h. Next, ovaries were washes 3 times in PBS-T for 10 min and washed once in PBS. Ovarioles were separated, mounted in ProLong™ Diamond Antifade Mountant (Thermo Fisher Scientific; P36961) and imaged 24 h later in a Zeiss LSM 880 confocal microscope (Carl Zeiss MicroImaging) with a Nikon EC Plan-Neofluar 40x/1.30 Oil DIC M27 objective.

### Correlative light and Electron Microscopy

*Drosophila* ovaries expressing GFP-Clu were fixed with 4 % formaldehyde (EM grade, Electron Microscopy Sciences; 15710) and 2.5 % glutaraldehyde (EM grade; Sigma-Aldrich; G5882) in PBS for one h at room temperature, and then overnight at 4 °C. Next, they were washed three times in PBS. Ovarioles were transferred to a 35 mm dish (MatTek, P35GC-1.5-14-C) and immobilised with a small drop of low melting point agarose with coverslip (No. 1.5), left to set and then submerged in PBS.

Confocal images were captured using an Olympus FV1000 confocal laser scanning microscope equipped with an UPlanSApo 60x/1.35 NA oil objective and zoom x3.5. Optical sections with a thickness of 250 nm were taken and the location and depth from the coverslip to positive labelled structures recorded. Deconvolution processing was performed using Huygens Essential version 21.04 (Scientific Volume Imaging, The Netherlands) with “Standard” profile settings. Following confocal imaging, samples were washed in 0.1 M cacodylate buffer for one h, and post fixed in 1 % osmium tetroxide/1.5 % potassium ferricyanide in 0.1 M cacodylate buffer for 1h. After further cacodylate washes, samples were stained with 1% tannic acid in 0.05 M cacodylate, followed by 5 min ‘stop’ in 1 % sodium sulphate in 0.05 M cacodylate buffer. Samples were washed in water, and serially dehydrated through a 70, 90 & 100 % ethanol series then propylene oxide, and finally flat embedded in TAAB 812 resin. Resin was polymerised at 60 °C for 48 h. 400 nm thick sections were taken using a Reichert Ultracut E ultramicrotome to a depth of 4000 nm, and then 70 nm thick serial sections were cut and collected onto formvar coated slot grids. Sections were further contrasted with Reynold’s lead citrate for 5 min before imaging at 120 kV on a JEOL JEM-1400 transmission electron microscope (TEM) (JEOL UK) using a Xarosa digital camera with Radius software (EMSIS, Germany).

Low magnification TEM images (x1500) were roughly correlated to confocal images using characteristic features such as nuclei and cell walls to identify potential regions of interest. Each sequential section was imaged at the ROI at higher magnifications (2.5K and 5K) to perform CLEM. Optical sections were correlated to TEM images utilising characteristic features as fiducial markers using Fiji software [65] with TrakEM2 plugin [66]. All EM consumables sourced from Agar scientific ltd, UK. unless indicated otherwise.

### RNase treatment

The RNase treatment was adapted from [36]. Briefly, ovaries from 2 GFP-Clu day-old-flies, kept with males and fed with yeast paste (n=5) were dissected in Schneider’s media (Lonza; LZ04-351Q) containing 200 μg/mL insulin (Sigma, I5500) (SI), then permeabilised in a buffer (100 mM potassium phosphate pH 7.5 and 0.2 % Triton X-100) for 50 sec. Then ovaries were washed twice with SI and incubated either with SI only (control) or SI containing 300 μg/mL of RNase A (Ambion; AM2270) for 15 min. After washing ovaries twice with SI, samples were mounted in SI and immediately imaged in a Zeiss LSM 880 confocal microscope (Carl Zeiss MicroImaging) with a Nikon Plan-Apochromat 63x/1.40 NA oil immersion objective.

### Cycloheximide and Puromycin treatment

One-day-old GFP-Clu female flies were crossed with males for one day in propionic food. Ovaries were dissected in Schneider’s media and were either subjected to no treatment or treated with 100 µg/mL CHX (Sigma-Aldrich; C7698) or 50 µg/ml puromycin (MP Biomedicals; 11420802) for 10 mins. Then, 200 μg/mL insulin (Sigma, I5500) was added and incubated for 30 min. As an additional control, ovaries were maintained without any drug or insulin for 40 min. Ovaries were then fixed, mounted and imaged as described above (n > 3 flies for each condition).

### 1,6-hexanediol treatment

One day old female flies were kept with males with yeast paste for two days. Ovaries from 5 flies were dissected in SI and placed onto a 35 mm dish (MatTek, P35GC-1.5-14-C) with SI. 4% of 1,6-hexanediol (Sigma-Aldrich; 240117) was added and snapshots were taken every min up to 15 mins in a Zeiss LSM 880 confocal microscope (Carl Zeiss MicroImaging) with a Nikon Plan-Apochromat 63x/1.40 NA oil immersion objective.

### Yeast methods Strains and plasmids

Yeast strains, plasmids and primers used in this study are detailed in Table S3. The *CLU1* knockout strain was generated in the W303a parental strain through PCR-based gene disruption using a HYGMX4 cassette, as previously described [67]. Clu1-mCherry and Clu1-TEV-GFP was generated by integrating mCherry or TEV-GFP in the C-terminal of *CLU1* gene in the BY4741 strain using a PCR-based gene insertion as previously described [68]. BirA* was integrated into the BY4741 Clu1-GFP strain immediately following the GFP gene, and into the flanking regions of the His3 gene in BY4741, using the same PCR-based insertion approach. The mCherry cassette was amplified from pBS35 (Addgene plasmid; 83797), TEV-GFP from pFA6a-GFP(S65T) (Addgene plasmid; 41598) using a 5’ primer containing the TEV sequence, and the BirA* cassette from pYM28-BirA* (Addgene plasmid; 160290). pRS313-pTDH3-Su9-TagBFP was generated using a yeast recombination-based cloning approach, replacing the URA3 marker from pRS316-pTDH3-Su9-TagBFP (Addgene plasmid # 62383) by HIS3. HIS3 and the flanking regions which are homologous to the pRS316 were amplified from a pRS413 plasmid and transformed into yeast with the NdeI linearised pRS316-pTDH3-Su9-TagBFP. Transformants were selected on SC without His plates and counter selected on SC without URA. Next, the plasmid was extracted from yeast using the QIAprep Spin Miniprep Kit (Qiagen; 27104) following the manufacturer instructions with an additional lysing step: after the addition of buffer P1, cells were lysed using acid-washed glass beads (425-600 μm diameter, Sigma-Aldrich, G8772) in a Precellys 24 homogenizer (Bertin Technologies), at 6,500 rpm for 15 sec with 5 min on ice, repeated 5 times. The extracted DNA was then transformed into DH5α competent cells (Invitrogen; 18265017) and transformed colonies were selected on LB containing Amplicillin. The correct deletion of *CLU1*, the insertion of mCherry, TEV-GFP, BirA* and the replacement of URA3 by HIS3 in pRS316-pTDH3-Su9-TagBFP were verified by colony PCR using primers that amplify across the flanking gene regions and by sequencing the inserts. BiFC strains were obtained from Bioneer (Korea). *CLU1-VN* strain (Mata) was mated with all the other VN strains (Matalpha) – *SOD2*, *PYC2*, *RPL17B*, *TIF3*, *ACO1*, *SCP160*, and *TIF5* to generate diploids, which were then selected in SC without uracil and leucine (SC-Ura-Leu). pRS416-MET25-ffRFP-MDV1 and pRS416-MET25-ffRFP-CAF4 [27] were kindly given by Janet M Shaw. pRP2132, Ded1-mCherry [69] by Roy Parker. pAG415GPD/Dcp1-dsRed construction was previously described [70].

### Media and growth conditions

*Saccharomyces cerevisiae* yeast strains were grown on standard media, YPD or Synthetic Complete (SC) with all amino acids or without specific amino acids according to the plasmid selection. YPD contains 10 g/L yeast extract, 20 g/L peptone and 20 g/L dextrose. SC media includes 6.7 g/L of yeast nitrogen base with ammonium sulphate and without amino acids, supplemented with a complete amino acid mixture or, a drop out mixture according to the plasmid selection (concentrations of amino acid mixtures were based on manufacturer’s instructions) and a carbon source. Depending on the auxotrophic markers, media was further supplemented with increased concentrations of histidine, leucine, methionine, uracil or adenine, to prevent amino acid starvation in the late log phase, as previously described [71]. Unless indicated otherwise, the carbon source used in most experiments was glucose (2%). Other carbon sources used were ethanol (3%), glycerol (3%) or galactose (3%). All media reagents were sourced from Formedium. To induce expression of Mdv1 and Caf4 from pRS416-MET25-ffRFP-CAF4 in the BY4743 Clu1-GFP strain, cells were grown in SC without Uracil and Methionine (SC -URA-MET).

Unless otherwise specified, all cultures were initially grown as pre-cultures in glucose-containing SC media. These pre-cultures were then diluted back into the same media to synchronize cells in the log phase overnight. For media shifts, early log-phase cells were washed in SC media without any carbon source and resuspended in media containing a different carbon source or no carbon source for starvation experiments. All cultures were incubated in a shaking incubator set at 30 °C, 225 rpm.

The determination of the diauxic shift was performed by measuring glucose levels in the culture media using the Glucose (GO) Assay Kit (Sigma-Aldrich; GAGO20). The start of the PD phase was defined as the time when cells began regrowing and oxygen consumption increased. Oxygen consumption was measured using an Oroboros oxygraphy-2K high-resolution respirometer (OROBOROS Instruments). Oxygen consumption rate was calculated by subtracting non-mitochondrial respiration (calculated after addition of 2 µM antimycin A) from the total oxygen consumption rate. Data acquisition and analysis were carried out using DatLab software (OROBOROS Instruments).

### Microscopy

To immobilise live cells, 1% agarose solution (UltrapureTM Agarose, Invitrogen) in appropriate SC media was prepared. Agarose pads were created by pipetting 30 µl of heated agarose solution onto a slide in between two layers of sticky-tape. Then, a second slide was placed on top of the other and allowed to set. Right before imaging, the top slide was slowly slid off, resulting in an intact agarose pad attached to the bottom slide. Subsequently, 7 µl of cells were pipetted onto the pad and covered with a coverslip. Alternatively, cells were directly pipetted into a slide and imaged immediately. For yeast fixation, cultures were incubated in 4% formaldehyde (Thermo Fisher Scientific; 28908) solution containing 4 % (w/v) sucrose made in PBS for 15 min at 30°C. After fixation, cells were washed twice in 1.2 M sorbitol and 0.1 M potassium phosphate buffer. Cells were mounted and immobilised in ProLong™ Diamond Antifade Mountant (Thermo Fisher Scientific; P36961) and imaged 24 h later.

Images were acquired with a Zeiss LSM 880 confocal microscope (Carl Zeiss MicroImaging) with Nikon Plan-Apochromat 63x/1.40 NA oil immersion objective, with a step size of 0.35 µm for z-stacks. Images were analysed with Fiji’s Image J [65].

### Stress treatments

Mid-log cells grown in SC with 2% of glucose were subjected to the following treatments: heat shocked at 46 °C for 30 min in a shaking heat block or maintained at 30 °C as control; incubated with 1M potassium chloride for 30 min; with 3 mM hydrogen peroxide for 15 min; washed once with water and incubated in water for 10 min, or SC without any carbon source for 10 min; incubated with sodium azide 0.5% for 30 min; treated with vehicle or 30 µM CCCP (prepared in 100% ethanol) for 15 min (unless otherwise stated) by direct addition into the media.

For stress tests in SC galactose, cells were grown as described above. After 6 h of incubation in galactose, when cells restarted growing, they were subjected to the same stress treatments as described for glucose-grown cells. Additionally, a mixture of 10 µM oligomycin and 40 µM antimycin A (OA) was tested for 30 min. For CHX tests, 100 µg/ml CHX (Sigma-Aldrich; C7698) (dissolved in 100% ethanol) or ethanol, as control, was added to Clu1-GFP cells in early PD cells (8 h after reaching optical density 5 at 600 nm absorbance (OD_600_) in SC media with 2% glucose). Cells were imaged before and after adding CHX or ethanol. The same concentration of CHX or ethanol was also added to cells that were growing in SC galactose (3%) for 6 h (shifted from SC glucose media), followed by treatment with 30 µM CCCP for 15 min. Cells were fixed as described in the “Microscopy” section.

### 1,6-hexanediol treatment

Cells in the PD phase were treated with 10 µg/ml digitonin for 2 min with or without 5 or 10% 1,6-hexanediol for 5 min. They were immediately fixed and imaged as described in the “*Microscopy*” section.

### Proximity-dependent biotinylation identification (BioID)

Pre-cultures of BirA* and Clu1-GFP-BirA* strains were established in quadruplicate in SC-His media to synchronize growth. For mid-log phase, early log phase cultures were diluted to reach an OD_600_ of 2.5 after 16 h of incubation with 50 μM biotin. For PD phase (PD), 50 μM biotin was added 8 h after reaching OD_600_ 5, and incubated for 16 h. After biotin incubation, cells were centrifuged (4,000 rpm, 5 min), washed in sterile water and flash frozen in liquid nitrogen. Cellswere lysed as previously described [72]. Briefly, 10% trichloroacetic acid was added to frozen cells (60 OD_600_), incubated on ice for 20 min and subsequently centrifuged 3 min at 15,000 g, at 4 °C. The pellet was washed with ice-cold acetone twice, dried at room temperature and then resuspended with 750 µl of MURB buffer (50 mM sodium phosphate, 25 mM MES, pH 7.0, 1% SDS, 3 M urea, 0.5% 2-mercaptoethanol, 1 mM sodium azide). Next, the pellet was disrupted by vortex with acid-washed glass beads (425-600 μm diameter, Sigma-Aldrich, G8772) in Precellys 24 homogenizer (Bertin Technologies), at 6,500 rpm for 15 sec with 5 min on ice, repeated 5 times. Next, the lysate was centrifuged at 18,000 g at 4 °C for 15 min and the supernatant transferred into a new tube.

For affinity purification, 600 µl Pierce^TM^ high-capacity streptavidin agarose Resins beads (Thermo Fisher Scientific, 20359) were first equilibrated by washing three times with affinity purification (AP) buffer (8 M Urea, 1 % SDS, 50 mM Tris-HCL Buffer (pH 7.4), 1 mM DTT, cOmplete™ EDTA-free Protease Inhibitor Cocktail (Roche)). After equilibration, the cell lysate was diluted to 7.5 ml with the AP buffer. Lysates were incubated with the equilibrated beads overnight, at room temperature. The following day, the beads were washed 5 times by centrifugation at 1,000 g for 2 min and resuspending beads in wash buffer (8 M Urea, 1 % SDS, 50 mM Tris-HCL Buffer (pH 7.4), 1 mM DTT). Biotinylated proteins were eluted in 750 µl Laemmli Sample Buffer (Bio-Rad, 1610747) containing β-mercaptoethanol (Sigma-Aldrich; M6250) incubating it at 95 °C for 10 min. Beads were removed from eluates using Vivaclear 0.8 µm PES centrifugal microfilters (Sartorius, VK01P042) and eluates were concentrated via Vivaspin 500 centrifugal concentrators (Merck, Z614068) with molecular weight cut-off of 30 kDa to remove free streptavidin. Proteins were resolved by SDS-PAGE and 6 gel regions from each sample were excised. Regions of gel containing a prominent 250 kDa protein, present in every sample, were excluded from the analyses.

Proteins were trypsinised within the gel. The tryptic peptide mixtures were desalted and purified by C18-reverse phase chromatography (ZipTip, Millipore) and then analysed by LC-MS/MS using a Proxeon EASY-nanoLC system, with a C18 column (50 µm x 150 mm) coupled directly to a Q-Exactive plus Orbitrap mass spectrometer (Thermo Fisher Scientific). Peptides were fragmented by collision induced dissociation with nitrogen. Raw MS data were analysed using MaxQuant software [73] as described in [74]. Data was searched against a sequence database of *Saccharomyces cerevisiae* proteins (UniProt) that was modified to include streptavidin sequence. Relative protein abundances were compared by label-free quantification. Data normalization and statistical analysis (t-test) were performed in Perseus software [75].

### Puromycin-associated nascent chain proteomics (Punch-P)

Punch-P was performed as described in [47, 48, 74]. WT and *clu1Δ* cells in the log phase were shifted from SC + glucose to SC + ethanol 3% and grown for 4 h (n=4). Yeast cells equivalent to 300 OD_600_ per condition were pelleted, washed once with cold water, snap-frozen, and stored at −70 °C. Each cell pellet was thawed and resuspended in 900 μl of polysome buffer (PLB) (20 mM Tris-HCl, pH 7.4, 140 mM KCl, 10 mM MgCl_2_, 0.5 mM DTT, 40 U/mL RNasin (Promega, N2111), 1.4 μg/mL pepstatin (Sigma-Aldric 10253286001), 2 μg/mL leupeptin (Sigma-Aldrich, 11017101001), 0.2 mg/mL heparin (Sigma-Aldrich), EDTA-free cOmplete protease inhibitor mix (Roche; 4693159001)) containing 1% Triton X-100. Cells were lysed using acid-washed glass beads (425-600 μm diameter, Sigma-Aldrich; G8772) corresponding to an approximate volume of 300 µL and homogenized in Precellys 24 (Bertin Technologies) at 6,500 rpm for 10 sec with 5 min on ice in between, repeated 5 times. Cell debris was removed by centrifuging the lysates at 17,400 g for 30 min at 4 °C. Next, 670 µL of lysate was slowly dispensed on top of 330 µL of a 70% sucrose cushion in a 1 mL polycarbonate tube (Beckman Coulter, 343778) and centrifuged in a MLA-130 ultracentrifuge rotor (Beckman Coulter) at 48,600 g for 160 min at 4 °C. The ribosome pellets were gently washed in 500 μl of ice-cold RNase-free water to remove all sucrose. Polysomes were carefully resuspended in 90 µl PLB (without detergent) until a homogeneous, slightly opaque solution is obtained. rRNA absorbance was measured at 254 nm and the volume corresponding to 15 OD_600_ was incubated with 1.5 nmol of Biotin-dC-puromycin (Jena Bioscience, NU-925-BIO-S) at 37 °C for 15 min to label nascent peptides. The same volume of polysomes was kept aside and used as control to identify endogenously biotinylated proteins and other contaminants (unlabelled). Labelled and unlabelled samples were incubated with 75 μl of Pierce™ Streptavidin Agarose (Thermo Fisher Scientific, 20347) in 1 mL of SDS/urea buffer (50 mM Tris-HCl, pH 7.5, 8 M urea, 2% SDS, and 200 mM NaCl) overnight at room temperature. Next day, beads were washed five times with SDS/urea buffer and incubated in 1 M NaCl for 30 min, followed by five washes in ultrapure water, 30 min incubation with 1 mM DTT, 30 min incubation with 50 mM iodoacetamide and two washes with 50 mM ammonium bicarbonate. Washes were achieved by centrifugation at 1,000 g for 2 min. Proteins were trypsinized on beads and identified by LC-MS/MS analysis as described in the “BioID” section. Proteins that were more abundant in both WT and *clu1Δ* unlabelled samples were considered as contaminants.

### Clu1 RNA immunoprecipitation (RIP)

Clu1-GFP and cells expressing GFP via p413-GPD-GFP were grown until the mid-log phase; cells equivalent to 50 OD_600_ were pelleted, flash frozen and stored at -70 °C. Cells were resuspended in 800 µL of PLB (20 mM HEPES pH 7, 150 mM KCl, 5 mM MgCl_2_, 0.5% NP-40, 1 mM DTT, 1 mM PMSF, 400 U/mL RNaseOUT (Thermo Fisher Scientific; 10777019) 100 µg/mL CHX (Sigma-Aldrich; C7698-1G), EDTA-free cOmplete protease inhibitor mix (Roche; 4693159001)) and lysed in Precellys 24 (Bertin Technologies) as described in the “*Punch-P”* section. For puromycin and EDTA treated samples, CHX was omitted from PLB. For EDTA treated samples, EDTA was added to lysates at 50 mM concentration. After lysis, cells were centrifuged at 3,000 rpm twice for 10 min at 4 °C. A 25 µl aliquot of the lysate was kept for RNA extraction (input). Lysates were pre-cleared with Dynabeads™ protein G beads (Thermo Scientific; 10003D) for 30 min, and then incubated with the beads conjugated with a mouse monoclonal GFP antibody (Abcam; ab1218) for 2 h 30 min at 4 °C, using an end over end rotator. Next, beads were washed 6 times with wash buffer (20 mM HEPES pH 7, 350 mM KCl, 5 mM MgCl_2_, 1%, NP-40, 0.1 mM DTT, 100 µg/mL CHX). CHX was not added to the wash buffer of samples treated with EDTA or puromycin. For puromycin treated samples, after the last wash, beads were resuspended with PLB containing 500 mM of KCl instead of 350 mM and 1 mM puromycin and incubated for 10 min at 37 °C. All samples were then resuspended with wash buffer containing 10 U of DNase I (Thermo Scientific; AM2235), incubated for 2 min at 37°C, and washed one more time with wash buffer. Finally, beads were resuspended in Tri reagent LS (Sigma-Aldrich; AM9738) to extract mRNAs. All buffer reagents were purchased as Molecular Biology, Nuclease-Free grade.

### Real time quantitative PCR (RT-qPCR)

RNA was extracted using Direct-zol RNA microprep kit (Zymo Research; R2060) following the manufacturer’s instructions. RNA extracted from polysome pellets (Punch-P) were treated with TURBO DNA-free™ Kit (Thermo Fisher Scientific; AM1907) to remove DNA contamination following the manufacturer’s instructions. cDNA was synthetised using Maxima H Minus cDNA Synthesis Kit with dsDNase (Thermo Fisher Scientific; M1681) following the manufacturer’s instructions. For polysome pellet samples, 500 ng of total RNA was used for cDNA synthesis, while for RNA extracted from RIP or input samples, the whole samples were used. cDNA was amplified using PowerUp™ SYBR™ Green Master Mix for qPCR (Thermo Fisher Scientific; A25741) following manufacturer’s instructions, in a QuantStudio™ 3 Real-Time PCR System (Applied Biosystems). Primers used are described in Table S3. Relative quantification was performed using the comparative CT method using PCR primer efficiency [76].

### Clu1 immunoprecipitation

Cells expressing Clu1-TEV-GFP or GFP (p413-GPD-GFP) were harvested in the log phase and immunopurified as described in the “RIP” section (PLB with CHX). After the last wash, an aliquot of beads was stored to run by WB (IP) and the remaining beads were incubated overnight at 4 °C with 100 μl PLB containing 0.5 μg of EDTA-free TEV protease (Sigma, T4455) to release Clu1 and co-interactors from the beads (eluate). Finally, the eluates were concentrated in Vivaspin 500 centrifugal concentrators with 10 kDa cut-off (Merck; Z614033). To elute proteins from IP samples, beads were incubated with 2x Laemmli Sample Buffer (Bio-Rad, 1610747) containing 1:10 β-mercaptoethanol (M6250; Sigma-Aldrich) for 10 min at 95 °C.

### Sucrose gradient ultracentrifugation

Clu1-GFP yeast cells were analysed in different growth conditions: 1) mid-log phase 2) PD 3) grown until early log in SC glucose, shifted to SC galactose and treated with CCCP as described in the “*Stress treatments”* section and 4) grown as in condition 3) and spiked with 2% glucose and incubated for 15 min. Cells were cross-linked with formaldehyde in all conditions, with exception of mid-log cells, which were also analysed without cross-linking. For cross-linking, media was removed by centrifugation at 4,000 rpm for 5 min and resuspended in 1% formaldehyde made in PBS and incubated with rotation for 20 min at room temperature. To quench the reaction, 125 mM of glycine was added for 5 min. Cells where then washed once with PBS, the pelleted cells were snap frozen with liquid nitrogen and stored at -70 °C. Cells equivalent to 50 OD_600_ were used for all conditions.

Frozen cell pellets were lysed in 600 μl of the same polysome buffer (PLB) described in the “*Clu1 RIP*” section, with the following modifications: 15 U of TURBO™ DNase (Thermo Fisher Scientific; AM2238) were added to PLB; for EDTA non-cross-linked treated samples, CHX was excluded from the buffer; for RNase treated samples, RNaseOUT was excluded from the buffer. Lysis was performed as described in the “*Punch-P”* section and debris was removed by spinning lysate twice at 3,000 rpm at 4 °C for 10 min. For EDTA non-cross-linked samples, 50 mM EDTA was added after lysis and for RNase treated samples, 1 μg of RNase A (Ambion; AM2270) was added per 600 μg of RNA and incubated for 30 min at room temperature. To stop the RNase A reaction, 200 U of RNaseOUT (Thermo Fisher Scientific; 10777019) were added for each μg of RNase A used. Sucrose continuous density gradients, 5-45% sucrose in PLB buffer (without RNaseOUT or DNase), were prepared using a Gradient Station (BioCOMP). For EDTA samples, 50 mM EDTA was also added to the buffers and CHX was excluded. Clarified lysates were layered on top of the gradients, ultracentrifuged in a SW 40 Ti rotor (Beckman Coulter) at 39,300 rpm, 4 °C, for 2 h 30 min, and fractionated using a Gradient Fractionator (Brandel). RNA profiling across gradients was performed by continuous monitoring of absorbance at 260 nm using an AKTA prime plus (Cytiva), which allowed identification of fractions enriched in ribosomal subunits, monosomes and polysomes (labelled in the immunoblots).

### Immunoblotting

Proteins were separated by SDS–PAGE in 4–20% Mini-PROTEAN® TGX™ Precast Protein Gels (Bio-Rad; 4561096) and then transferred onto a nitrocellulose membrane (Bio-Rad; 1704158) using the Trans-Blot Turbo Transfer System. Membranes were blocked for 1 h at room temperature with 5% (wt/vol) dried skimmed milk powder (Marvel Instant Milk) in TBS containing 0.1% Tween-20 (TBS-T). Following blocking, membranes were incubated overnight at 4°C with the appropriate primary antibodies diluted in TBS-T with exception of anti-Pgk1 which was only incubated for 1 h. After three washes with TBS-T for 10 min, membranes were incubated for 1 h at room temperature with HRP-conjugated secondary antibodies diluted in 5% milk in TBS-T. Membranes were then washed four times, for 10 min in TBS-T, and antibody detection was performed using the Amersham ECL Prime detection kit (Cytiva, RPN2232). Sucrose gradient immunoblots were stripped after incubation with anti-Rpl3 with Restore™ Western Blot Stripping Buffer (Thermo Fisher Scientific; 21059) according to manufacturer’s instructions before incubating with anti-Pgk1. The immunoblots were imaged with the Amersham Imager 680.

The primary antibodies used for immunoblotting in this study were: anti-GFP (1:1000; Abcam; ab290), anti-BirA (1:1000; Novus Biologicals; NBP2-59939), anti-Pgk1 (1:10000; Thermo Fisher Scientific; 459250) and anti-Rpl3 (1:2000; DHSB). Secondary antibodies used were: HRP-conjugated goat anti-mouse IgG H&L (1:20000; Abcam; ab6789) and HRP-conjugated goat anti-rabbit IgG H&L (1:10000; Thermo Fisher Scientific; G-21234).

### Yeast spotting assay

WT and *clu1Δ* strains were grown in SC containing glucose and diluted once back to synchronize cells to early log phase. Cell concentration was adjusted to be equal among strains (OD_600_ = 1) and 5-fold serial dilutions were prepared with water in a 96-well plate. Next, they were spotted onto SC containing 2% glucose or 3% ethanol plates. Plates were left in the 30 °C incubator until growth was observed. Images were obtained with a GelDoc XRS+ (BioRad).

### Statistical analysis

GraphPad Prism 9 (SCR_002798; RRID) was used to perform all statistical analyses. Two-samples groups were analysed by unpaired t-test with Welch’s correction. Multiple sample groups were compared using one-way ANOVA with Tukey’s post-hoc test.

## Declarations

### Data availability

All data needed to evaluate the conclusions in the paper are present in the paper and/or the Supplementary Materials.

## Acknowledgements

We kindly thank Rachel Cox (Uniformed Services University), Daniel St Johnston (University of Cambridge) and Erika Geisbrecht (Kansas State University) for generously sharing fly lines, and Nianshu Zhang (University of Cambridge), Roy Parker (University of Colorado Boulder), Janet M Shaw (University of Utah), Benjamin Glick (University of Chicago), Jonathan Weissman (Whitehead Institute), Eric Muller (University of Washington), John Pringle (Stanford University) and Martin Ott (University of Gothenburg) for kindly sharing yeast strains and plasmids. We thank Ian Fearnley and Shujing Ding (MRC Mitochondrial Biology Unit) for technical assistance with the mass spectrometry and Natalie Allcock, Anna Straatman-Iwanowska and Kees Straatman (University of Leicester) for the CLEM work. Finally, we thank Jeffrey Gerst (Weizman Institute of Science) for technical assistance, Brian Zid and Yuko Sugiyama (University of California, San Diego) and all members of the Whitworth lab for discussions during the project and feedback on the manuscript.

This work was supported by European Research Council Starting Grant (309742), Medical Research Council (MRC) core funding (MC_UU_00015/6 and MC_UU_0028/6), MRC project grants; MR/V003933/1, and Motor Neurone Disease Association grant (Whitworth/Apr17/857-79).

## Author contributions

LMF: Conceptualisation, Methodology, Formal analysis, Investigation, Writing – Original Draft, Writing – Review & Editing, Visualisation, Supervision, Funding acquisition

WHA: Conceptualisation, Methodology, Investigation, Writing – Review & Editing

LR: Methodology, Formal analysis, Investigation

PRG: Conceptualisation, Methodology, Formal analysis, Investigation, Writing – Review & Editing JS: Methodology, Formal analysis, Investigation

HYC: Methodology, Formal analysis, Investigation

AC: Methodology, Formal analysis

AJW: Conceptualisation, Validation, Formal analysis, Writing – Original Draft, Writing – Review & Editing, Supervision, Project administration, Funding acquisition

## Supplementary Data

**Figure S1. Clu/Clu1 has a punctate subcellular distribution in several fly tissues and yeast upon entry into the PD phase.**

(A) Confocal imaging of the indicated GFP-Clu third instar larval and 2-day-old adult tissues. VNC, ventral nerve cord. Scale bars: 40 µm (top panels) and 8 µm (bottom panels, magnified images of boxed areas). (B) Maximum intensity projection of an egg chamber of a 3-day-old female fly refed for 6 h. A region of interest is marked in the image (white box) indicates the region used for 3D rendering in (C). (D, E) Box plots representing the area (D) and volume (E) of Clu foci from the region used for 3D rendering. The plots display the minimum, first and third quartile, median and maximum values. (F) Graph showing Clu1-GFP cells’ growth in glucose containing media, percentage of cells containing Clu1-GFP foci and basal respiration throughout growth. Cells undergo a diauxic shift upon exhaustion of glucose from the media. During this time, cells cease growth and suffer a drastic transcriptomic and proteomic shift to adapt to the new carbon source available in the media, ethanol. They go from a fermentative to a respiratory metabolism. We considered the start of PD phase the moment cells start regrowing and respiration increased.

**Figure S2. *Drosophila* GFP-Clu does not colocalise with SG or PB markers.**

(A, B) Confocal microscopy of GFP-Clu egg chambers fasted (16 h) or refed (6 h) immunostained for (A) Fmr1 (SG marker) or (B) co-expressing Tral-mRFP (PB marker). Scale bars = 20 µm, inset 4 µm.

**Figure S3. Localisation of Clu1 and the effects of its absence on growth and mitochondrial morphology.**

(A) Confocal image of Clu1-GFP cells expressing mito-mCherry in the PD phase. (B) Confocal microscopy single plane images of PD yeast cells expressing Clu1-GFP and ffRFP-Caf4, a marker of mitochondrial fission, and the maximum intensity projection (max proj) comprising the whole cells. (C, D) Spotting assay of *clu1Δ* and the parental strain W303A. Mid-log phase cells grown in glucose-containing media were adjusted to the same optical density, five-fold serial diluted and spotted onto media containing either glucose (C) or ethanol (D) as carbon sources. (E, F) Clu1-GFP cells expressing mito-mCherry were imaged by confocal microscopy in media containing either glucose (E) or ethanol (F). Scale bars = 5 µm.

**Figure S4, related to** Figure 6**. Validation of Clu1-GFP-BirA* strain.**

(A) Immunoblot of BirA* and Clu1-GFP-BirA* strains grown in the log and PD phases. (B) Confocal images of Clu1-GFP and Clu1-GFP-BirA* cells in the log and PD phases. Scale bar = 5 µm.

**Table S1. Label-free quantification and hit summary of BioID analysis.**

**Table S2. Label-free quantification and hit summary of Punch-P analysis.**

**Video S1. Addition of glucose to Clu1-GFP cells in PD phase.**

Clu1-GFP cells in the PD phase were layered on an agarose pad made with media containing 2% glucose and imaged over a 30 min time course.

**Video S2. Addition of insulin to GFP-Clu egg chambers.**

Time course over 62 min, imaged at one-min intervals, shows dissected egg chambers from two-day-old GFP-Clu flies following the addition of insulin.

## References

1. Friedman, J.R. and J. Nunnari, Mitochondrial form and function. Nature, 2014. 505(7483): p. 335–43.

2. Monzel, A.S., J.A. Enriquez, and M. Picard, Multifaceted mitochondria: moving mitochondrial science beyond function and dysfunction. Nat Metab, 2023. 5(4): p. 546–562.

3. Tilokani, L., et al., Mitochondrial dynamics: overview of molecular mechanisms. Essays Biochem, 2018. 62(3): p. 341–360.

4. Mishra, P. and D.C. Chan, Metabolic regulation of mitochondrial dynamics. J Cell Biol, 2016. 212(4): p. 379–87.

5. Chen, H. and D.C. Chan, Mitochondrial dynamics--fusion, fission, movement, and mitophagy--in neurodegenerative diseases. Hum Mol Genet, 2009. 18(R2): p. R169–76.

6. Rath, S., et al., MitoCarta3.0: an updated mitochondrial proteome now with sub-organelle localization and pathway annotations. Nucleic Acids Res, 2021. 49(D1): p. D1541–D1547.

7. Graham, L.C., et al., Proteomic profiling of neuronal mitochondria reveals modulators of synaptic architecture. Mol Neurodegener, 2017. 12(1): p. 77.

8. Morgenstern, M., et al., Quantitative high-confidence human mitochondrial proteome and its dynamics in cellular context. Cell Metab, 2021. 33(12): p. 2464–2483 e18.

9. Tsuboi, T., J. Leff, and B.M. Zid, Post-transcriptional control of mitochondrial protein composition in changing environmental conditions. Biochem Soc Trans, 2020. 48(6): p. 2565–2578.

10. Schatton, D. and E.I. Rugarli, A concert of RNA-binding proteins coordinates mitochondrial function. Crit Rev Biochem Mol Biol, 2018. 53(6): p. 652–666.

11. Fields, S.D., M.N. Conrad, and M. Clarke, The S. cerevisiae CLU1 and D. discoideum cluA genes are functional homologues that influence mitochondrial morphology and distribution. J Cell Sci, 1998. 111 **(Pt** **12**): p. 1717–27.

12. Cox, R.T. and A.C. Spradling, Clueless, a conserved Drosophila gene required for mitochondrial subcellular localization, interacts genetically with parkin. Dis Model Mech, 2009. 2(9-10): p. 490–9.

13. Gao, J., et al., CLUH regulates mitochondrial biogenesis by binding mRNAs of nuclear-encoded mitochondrial proteins. J Cell Biol, 2014. 207(2): p. 213–23.

14. Zhu, Q., et al., The cluA-mutant of Dictyostelium identifies a novel class of proteins required for dispersion of mitochondria. Proc Natl Acad Sci U S A, 1997. 94(14): p. 7308–13.

15. Schatton, D., et al., CLUH regulates mitochondrial metabolism by controlling translation and decay of target mRNAs. J Cell Biol, 2017. 216(3): p. 675–693.

16. Sen, A. and R.T. Cox, Clueless is a conserved ribonucleoprotein that binds the ribosome at the mitochondrial outer membrane. Biol Open, 2016. 5(2): p. 195–203.

17. Pla-Martin, D., et al., CLUH granules coordinate translation of mitochondrial proteins with mTORC1 signaling and mitophagy. EMBO J, 2020. 39(9): p. e102731.

18. Sheard, K.M., et al., Clueless forms dynamic, insulin-responsive bliss particles sensitive to stress. Dev Biol, 2020. 459(2): p. 149–160.

19. Buchan, J.R., D. Muhlrad, and R. Parker, P bodies promote stress granule assembly in Saccharomyces cerevisiae. J Cell Biol, 2008. 183(3): p. 441–55.

20. Buchan, J.R., J.-H. Yoon, and R. Parker, Stress-specific composition, assembly and kinetics of stress granules in Saccharomyces cerevisiae. Journal of Cell Science, 2011. 124(2): p. 228–239.

21. Shah, K.H., et al., Processing body and stress granule assembly occur by independent and differentially regulated pathways in Saccharomyces cerevisiae. Genetics, 2013. 193(1): p. 109–23.

22. Shah, K.H., et al., Processing Body and Stress Granule Assembly Occur by Independent and Differentially Regulated Pathways in Saccharomyces cerevisiae. Genetics, 2013. 193(1): p. 109–123.

23. Teixeira, D., et al., Processing bodies require RNA for assembly and contain nontranslating mRNAs. Rna, 2005. 11(4): p. 371–82.

24. Buchan, J.R. and R. Parker, Eukaryotic stress granules: the ins and outs of translation. Mol Cell, 2009. 36(6): p. 932–41.

25. Jain, S., et al., ATPase-Modulated Stress Granules Contain a Diverse Proteome and Substructure. Cell, 2016. 164(3): p. 487–98.

26. Rao, B.S. and R. Parker, Numerous interactions act redundantly to assemble a tunable size of P bodies in Saccharomyces cerevisiae. Proceedings of the National Academy of Sciences, 2017. 114(45): p. E9569–E9578.

27. Guo, Q., et al., The mitochondrial fission adaptors Caf4 and Mdv1 are not functionally equivalent. PLoS One, 2012. 7(12): p. e53523.

28. Lin, Y., et al., Formation and Maturation of Phase-Separated Liquid Droplets by RNA-Binding Proteins. Molecular Cell, 2015. 60(2): p. 208–219.

29. Patel, S.S., et al., Natively unfolded nucleoporins gate protein diffusion across the nuclear pore complex. Cell, 2007. 129(1): p. 83–96.

30. Kroschwald, S., et al., Promiscuous interactions and protein disaggregases determine the material state of stress-inducible RNP granules. eLife, 2015. 4: p. e06807.

31. Kroschwald, S., S. Maharana, and A.W. Simon. Hexanediol: a chemical probe to investigate the material properties of membrane-less compartments. 2017.

32. Itoh, Y., et al., 1,6-hexanediol rapidly immobilizes and condenses chromatin in living human cells. Life Sci Alliance, 2021. 4(4).

33. Bevilacqua, P.C., et al., RNA multimerization as an organizing force for liquid-liquid phase separation. Rna, 2022. 28(1): p. 16–26.

34. Garcia-Jove Navarro, M., et al., RNA is a critical element for the sizing and the composition of phase-separated RNA-protein condensates. Nat Commun, 2019. 10(1): p. 3230.

35. van Leeuwen, W. and C. Rabouille, Cellular stress leads to the formation of membraneless stress assemblies in eukaryotic cells. Traffic, 2019. 20(9): p. 623–638.

36. Lin, M.D., et al., Drosophila processing bodies in oogenesis. Dev Biol, 2008. 322(2): p. 276–88.

37. Kedersha, N., et al., Stress granules and processing bodies are dynamically linked sites of mRNP remodeling. J Cell Biol, 2005. 169(6): p. 871–84.

38. Eulalio, A., et al., P-body formation is a consequence, not the cause, of RNA-mediated gene silencing. Mol Cell Biol, 2007. 27(11): p. 3970–81.

39. Blobel, G. and D. Sabatini, Dissociation of mammalian polyribosomes into subunits by puromycin. Proc Natl Acad Sci U S A, 1971. 68(2): p. 390–4.

40. Sears, R.M., D.G. May, and K.J. Roux, BioID as a Tool for Protein-Proximity Labeling in Living Cells. Methods Mol Biol, 2019. 2012: p. 299–313.

41. Kim, D.I., et al., Probing nuclear pore complex architecture with proximity-dependent biotinylation. Proc Natl Acad Sci U S A, 2014. 111(24): p. E2453–61.

42. Hibbs, M.A., et al., Exploring the functional landscape of gene expression: directed search of large microarray compendia. Bioinformatics, 2007. 23(20): p. 2692–9.

43. Sung, M.K. and W.K. Huh, Bimolecular fluorescence complementation analysis system for in vivo detection of protein-protein interaction in Saccharomyces cerevisiae. Yeast, 2007. 24(9): p. 767–75.

44. Hu, C.D. and T.K. Kerppola, Simultaneous visualization of multiple protein interactions in living cells using multicolor fluorescence complementation analysis. Nat Biotechnol, 2003. 21(5): p. 539–45.

45. Sung, M.K., et al., Genome-wide bimolecular fluorescence complementation analysis of SUMO interactome in yeast. Genome Res, 2013. 23(4): p. 736–46.

46. Kim, Y., et al., Global analysis of protein homomerization in Saccharomyces cerevisiae. Genome Res, 2019. 29(1): p. 135–145.

47. Segev, N. and J.E. Gerst, Specialized ribosomes and specific ribosomal protein paralogs control translation of mitochondrial proteins. J Cell Biol, 2018. 217(1): p. 117–126.

48. Aviner, R., T. Geiger, and O. Elroy-Stein, PUNCH-P for global translatome profiling: Methodology, insights and comparison to other techniques. Translation (Austin), 2013. 1(2): p. e27516.

49. Young, J.C., N.J. Hoogenraad, and F.U. Hartl, Molecular chaperones Hsp90 and Hsp70 deliver preproteins to the mitochondrial import receptor Tom70. Cell, 2003. 112(1): p. 41–50.

50. Watson, H.C., et al., Sequence and structure of yeast phosphoglycerate kinase. The EMBO Journal, 1982. 1(12): p. 1635–1640.

51. Zaninello, M., et al., CLUH maintains functional mitochondria and translation in motoneuronal axons and prevents peripheral neuropathy. Sci Adv, 2024. 10(22): p. eadn2050.

52. Cho, E., et al., Cluh plays a pivotal role during adipogenesis by regulating the activity of mitochondria. Sci Rep, 2019. 9(1): p. 6820.

53. Sen, A. and R.T. Cox, Loss of Drosophila Clueless differentially affects the mitochondrial proteome compared to loss of Sod2 and Pink1. Front Physiol, 2022. 13: p. 1004099.

54. Ma, J., et al., Friendly mediates membrane depolarization-induced mitophagy in Arabidopsis. Curr Biol, 2021. 31(9): p. 1931–1944.e4.

55. Sieber, M.H. and A.C. Spradling, The role of metabolic states in development and disease. Curr Opin Genet Dev, 2017. 45: p. 58–68.

56. Sieber, M.H., M.B. Thomsen, and A.C. Spradling, Electron Transport Chain Remodeling by GSK3 during Oogenesis Connects Nutrient State to Reproduction. Cell, 2016. 164(3): p. 420–32.

57. Di Bartolomeo, F., et al., Absolute yeast mitochondrial proteome quantification reveals trade-off between biosynthesis and energy generation during diauxic shift. Proc Natl Acad Sci U S A, 2020. 117(13): p. 7524–7535.

58. Banani, S.F., et al., Biomolecular condensates: organizers of cellular biochemistry. Nat Rev Mol Cell Biol, 2017. 18(5): p. 285–298.

59. McSwiggen, D.T., et al., Evaluating phase separation in live cells: diagnosis, caveats, and functional consequences. Genes Dev, 2019. 33(23-24): p. 1619–1634.

60. Wheeler, J.R., et al., Distinct stages in stress granule assembly and disassembly. Elife, 2016. 5.

61. Hemono, M., et al., The interactome of CLUH reveals its association to SPAG5 and its co-translational proximity to mitochondrial proteins. BMC Biol, 2022. 20(1): p. 13.

62. Hemono, M., et al., FRIENDLY (FMT) is an RNA binding protein associated with cytosolic ribosomes at the mitochondrial surface. Plant J, 2022. 112(2): p. 309–321.

63. Nedelsky, N.B. and J.P. Taylor, Bridging biophysics and neurology: aberrant phase transitions in neurodegenerative disease. Nat Rev Neurol, 2019. 15(5): p. 272–286.

64. Lowe, N., et al., Analysis of the expression patterns, subcellular localisations and interaction partners of Drosophila proteins using a pigP protein trap library. Development, 2014. 141(20): p. 3994–4005.

65. Schindelin, J., et al., Fiji: an open-source platform for biological-image analysis. Nat Methods, 2012. 9(7): p. 676–82.

66. Cardona, A., et al., TrakEM2 software for neural circuit reconstruction. PLoS One, 2012. 7(6): p. e38011.

67. Wach, A., PCR-synthesis of marker cassettes with long flanking homology regions for gene disruptions in S. cerevisiae. Yeast, 1996. 12(3): p. 259–65.

68. Janke, C., et al., A versatile toolbox for PCR-based tagging of yeast genes: new fluorescent proteins, more markers and promoter substitution cassettes. Yeast, 2004. 21(11): p. 947–62.

69. Hilliker, A., et al., The DEAD-box protein Ded1 modulates translation by the formation and resolution of an eIF4F-mRNA complex. Mol Cell, 2011. 43(6): p. 962–72.

70. Miller-Fleming, L., et al., Yeast DJ-1 superfamily members are required for diauxic-shift reprogramming and cell survival in stationary phase. Proc Natl Acad Sci U S A, 2014. 111(19): p. 7012–7.

71. Murakami, C.J., et al., Composition and acidification of the culture medium influences chronological aging similarly in vineyard and laboratory yeast. PLoS One, 2011. 6(9): p. e24530.

72. Miller-Fleming, L., et al., Detection of Saccharomyces cerevisiae Atg13 by western blot. Autophagy, 2014. 10(3): p. 514–7.

73. Tyanova, S., et al., Visualization of LC-MS/MS proteomics data in MaxQuant. Proteomics, 2015. 15(8): p. 1453–6.

74. Aviner, R., T. Geiger, and O. Elroy-Stein, Genome-wide identification and quantification of protein synthesis in cultured cells and whole tissues by puromycin-associated nascent chain proteomics (PUNCH-P). Nat Protoc, 2014. 9(4): p. 751–60.

75. Tyanova, S., et al., The Perseus computational platform for comprehensive analysis of (prote)omics data. Nat Methods, 2016. 13(9): p. 731–40.

76. Pfaffl, M.W., A new mathematical model for relative quantification in real-time RT-PCR. Nucleic Acids Res, 2001. 29(9): p. e45.

