## Supplementary figures and images for "Clu1/Clu form mitochondria-associated granules upon metabolic transitions and regulate mitochondrial protein translation via ribosome interactions"

### Supplementary Figure S1

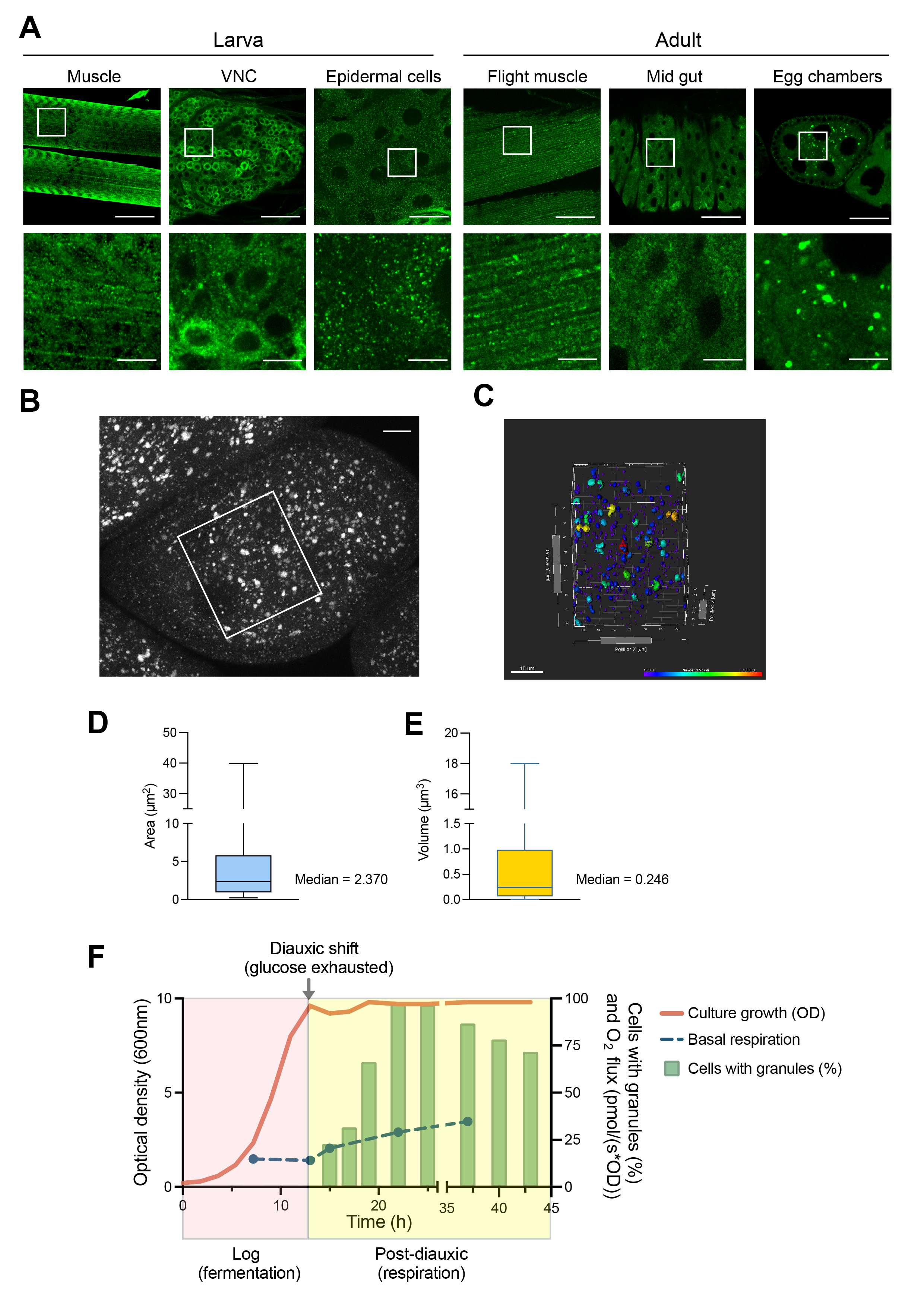

### Supplementary Figure S2

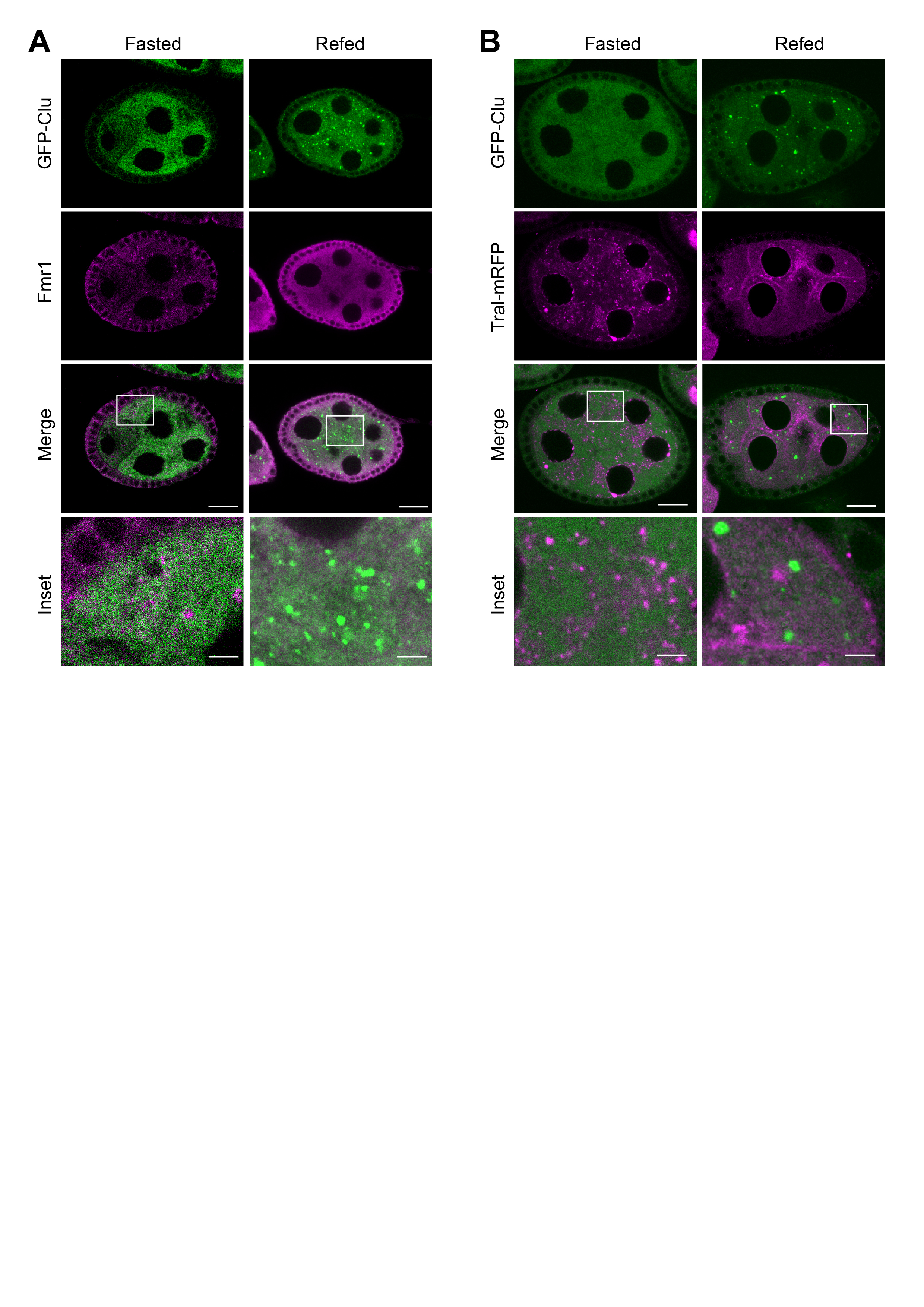

### Supplementary Figure S3

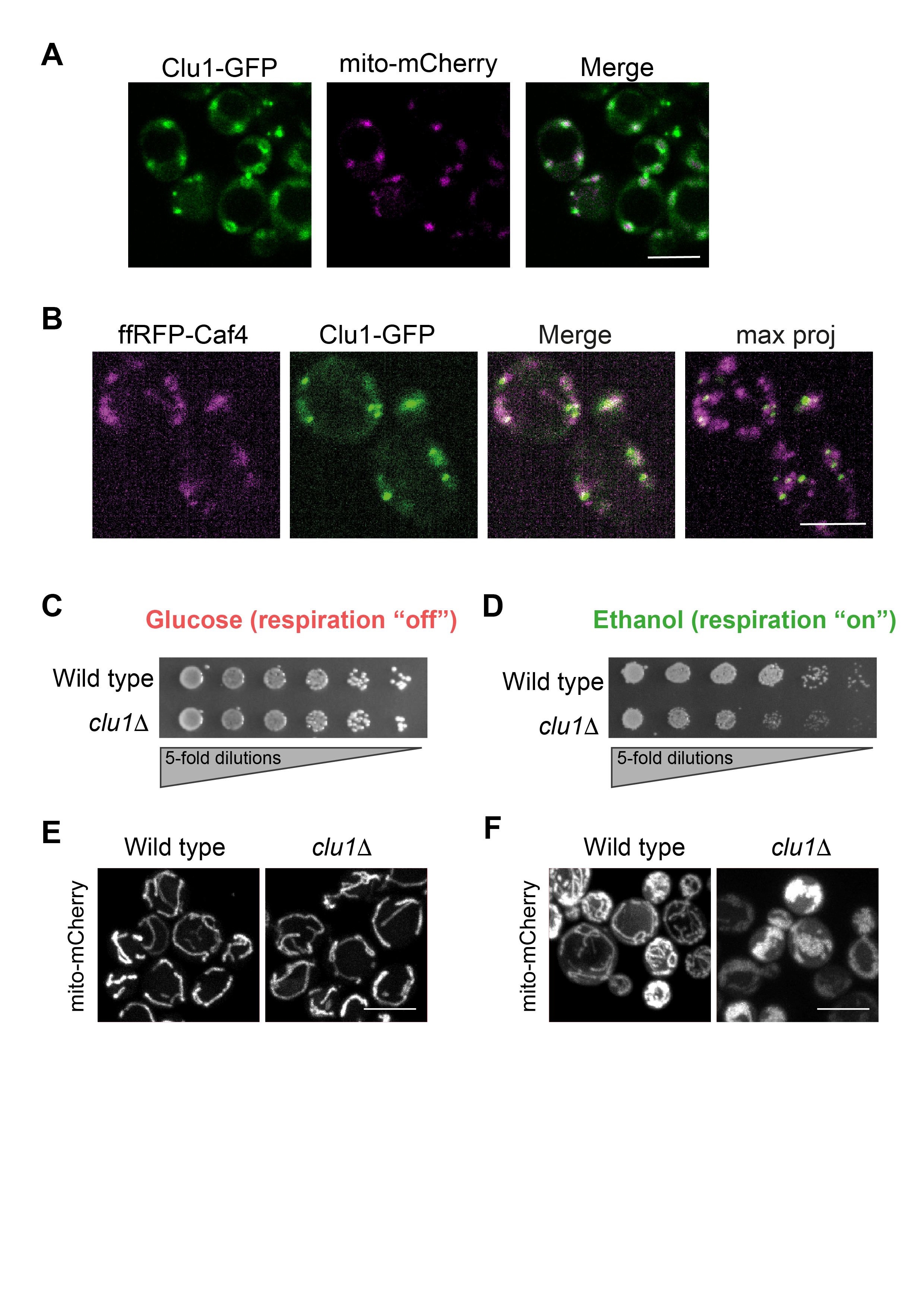

### Supplementary Figure S4

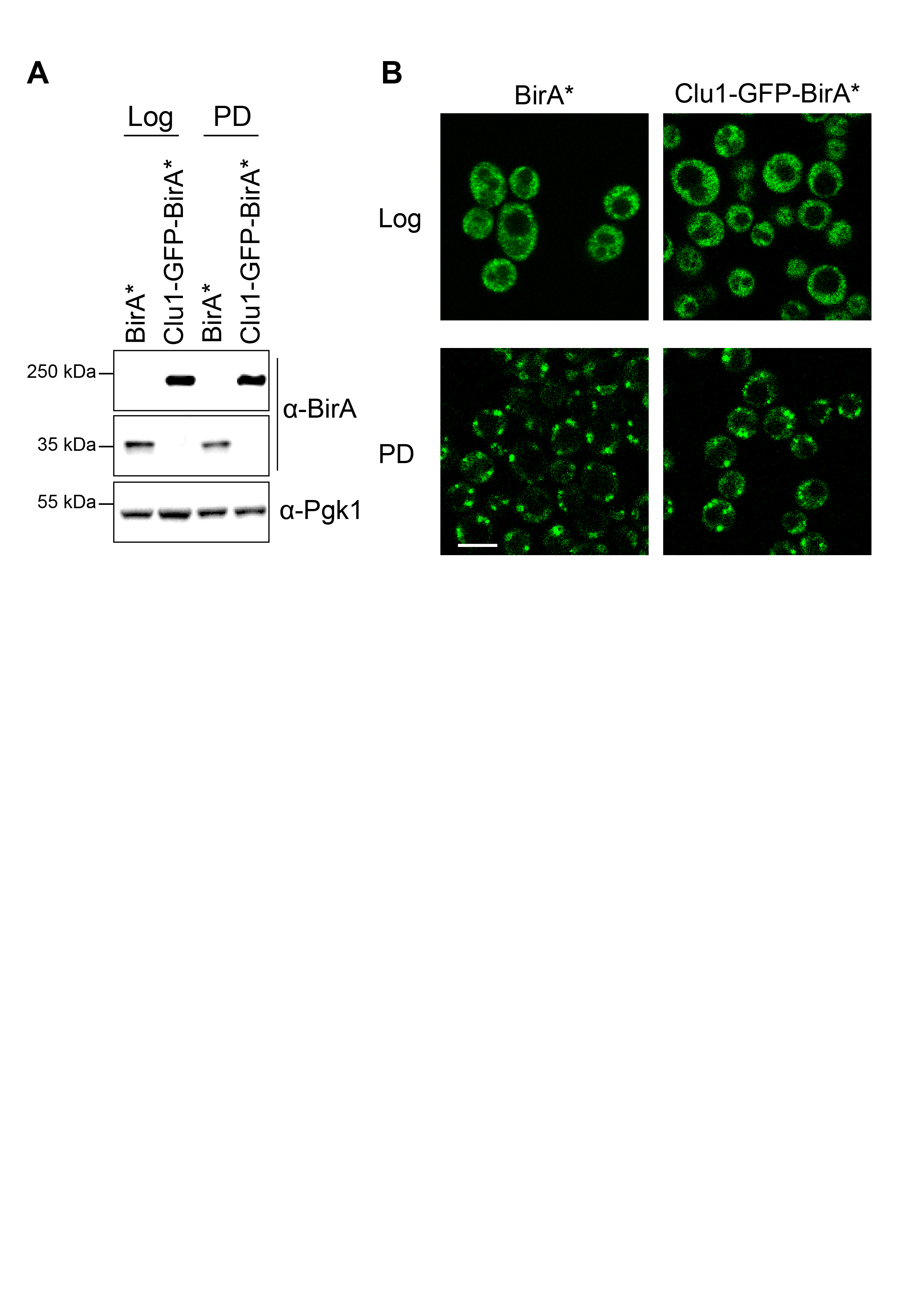
